# Retrosplenial cortex vulnerability links severe hypoglycemia to cognitive impairment through neuron–microglia crosstalk

**DOI:** 10.64898/2026.03.27.714654

**Authors:** Jae-Young Joo, Soyun Lee, Min Kyoung Shin, Sunmin Kim, Sohee Park, Jun H. Heo, Minji Kim, Hyojung Lee, Kyunghyuk Park, Dongjun Koo, Ho-Young Lee, Jong-Il Kim, Obin Kwon

**Affiliations:** Department of Biomedical Sciences, Seoul National University College of Medicine; 103 Daehak-ro, Jongno-gu, Seoul, Republic of Korea; Department of Biochemistry and Molecular Biology, Seoul National University College of Medicine; 103 Daehak-ro, Jongno-gu, Seoul, Republic of Korea; Genomic Medicine Institute (GMI), Medical Research Center, Seoul National University; 103 Daehak-ro, Jongno-gu, Seoul, Republic of Korea; Interdisciplinary Program in Bioengineering, Seoul National University; 1 Gwanak-ro, Seoul, Republic of Korea; Department of Nuclear Medicine, Seoul National University Bundang Hospital; 172 Dolma-ro, Seongnam-si, Gyeonggi-do, Republic of Korea; Department of Nuclear Medicine, Seoul National University College of Medicine; 103 Daehak-ro, Jongno-gu, Seoul, Republic of Korea; Convergence Dementia Research Center, Medical Research Center, Seoul National University; 103 Daehak-ro, Jongno-gu, Seoul, Republic of Korea

## Abstract

Severe hypoglycemia remains a serious adverse effect of insulin therapy in individuals with diabetes and is linked to cognitive decline, yet the mechanisms by which transient metabolic stress leads to persistent neuronal dysfunction remain poorly defined. Using mouse models of acute severe hypoglycemia and integrated screening, we identified the retrosplenial cortex as a previously unrecognized brain region that is particularly vulnerable to hypoglycemia-induced neuronal damage. This injury is driven by a feedforward interaction between neuron-specific Drp1-dependent mitochondrial fission and microglial IL-1 signaling, as pharmacological or genetic targeting of either pathway suppressed the other, rescued neuronal damage, and reversed cognitive impairment. These findings identify a region-specific neuron-microglia injury circuit that links severe hypoglycemia to cognitive dysfunction and suggest a therapeutic strategy to protect brain function without compromising diabetes management.

## INTRODUCTION

Maintaining optimal blood glucose levels is a cornerstone of diabetes management, with intensive glycemic control demonstrated to reduce the long-term risk of diabetic complications. To achieve this, insulin-based therapies have markedly improved glycemic management and patient outcomes^1^. However, insulin therapy carries the serious clinical risk of severe hypoglycemia, a condition characterized by critically low blood glucose levels requiring external assistance. Severe hypoglycemia is one of the most dangerous adverse effects of diabetes treatment and can lead to substantial brain injury^2,3^. Clinical studies have reported cognitive impairment following severe hypoglycemia^4^. Even with sensor-augmented insulin pump therapy, patients remain at risk of sudden death due to severe hypoglycemia^5^. As insulin requirements for diabetes management are projected to increase^6^, and hospitalization for severe hypoglycemia continues to impose a growing healthcare burden^7^, this clinical challenge is likely to become increasingly significant. Although intensive insulin treatment is associated with improved long-term outcomes^1^, the risk of severe hypoglycemia limits its broader application. Critically, no targeted neuroprotective strategy is currently available to prevent hypoglycemia-induced brain injury, largely because the brain regions selectively vulnerable to hypoglycemic insult and the cell type-specific mechanisms driving this damage remain poorly defined.

Because the brain is metabolically heterogeneous, severe hypoglycemia may not affect all regions or cell types equally. Identifying the neural circuits and cellular programs that are selectively vulnerable to hypoglycemic stress is therefore an important step toward mechanism-based neuroprotection. Among the candidate processes, mitochondrial homeostasis is of particular interest because mitochondrial fission and fusion are tightly coupled to cellular adaptation under metabolic challenge^8–10^. However, it remains unclear whether severe hypoglycemia induces region- and cell type-specific alterations in mitochondrial dynamics, and whether such changes contribute directly to neuronal injury.

Neuroinflammation may represent an important determinant of how severe hypoglycemia is translated into brain injury. In the injured brain, activated microglia release proinflammatory cytokines, including interleukin-1β (IL-1β), interleukin-6 (IL-6), and tumor necrosis factor-α (TNF-α)^11,12^. Severe hypoglycemia has been associated with microglial activation^13^, but it remains unclear which inflammatory mediators are most relevant to hypoglycemia-induced neuronal injury, whether they operate within selectively vulnerable brain regions, and how they interact with neuronal stress pathways to drive persistent dysfunction.

In this study, we identified the retrosplenial cortex as a selectively vulnerable brain region after severe hypoglycemia, with associated spatial memory impairment in mouse models. Our findings suggest that this damage is driven by a bidirectional interaction between neuronal Drp1-dependent mitochondrial fission and microglial activation, mediated in part by IL-1 signaling. Targeting this neuron-microglia circuit attenuated neuronal damage and mitigated cognitive impairment, supporting a potential therapeutic strategy to protect brain function without compromising peripheral diabetes management.

## RESULTS

### Oxidative stress, apoptosis, and neuronal dendritic loss in the RSC develop progressively after acute hypoglycemia

Previous studies in humans and rodents have reported that the cerebral cortex and hippocampus are more vulnerable than other brain regions to hypoglycemia-induced brain damage^14,15^. Thus, our initial aim was to identify the specific regions in the cortex and hippocampus that are vulnerable to acute hypoglycemia. To induce severe hypoglycemia, mice were fasted for 24 hours, and insulin or saline was intraperitoneally injected to maintain blood glucose levels below 20 mg/dL for 5 hours (Fig. 1a,b). Seven days after acute hypoglycemia induction, we conducted an unbiased screening via immunohistochemical staining for 4-hydroxynonenal (4-HNE), an oxidative damage marker, across the cortex and the hippocampus (dentate gyrus (DG) and CA1 and CA3 regions) (Supplementary Fig. 1a-i). Notably, an increased number of 4-HNE^+^ cells was observed in the retrosplenial cortex (RSC) region (Fig. 1c), whereas in other cortical regions (motor, somatosensory, visual, and auditory cortex), 4-HNE^+^ cells were absent or were present in comparable numbers between the hypoglycemia group and the control group (Supplementary Fig. 1b,c). Furthermore, the mean fluorescence intensity of 4-HNE in the hippocampus was comparable between the hypoglycemia group and the control group (Supplementary Fig. 1d-i). Importantly, [^18^F]-FDG PET/CT analysis demonstrated that the RSC also exhibited a significant reduction in regional cerebral glucose metabolism in hypoglycemia-experienced mice compared with controls (p < 0.01; Fig. 1d). These findings thus suggest that among the cortical and hippocampal regions, the RSC is particularly vulnerable to acute hypoglycemia.

**Fig. 1.**
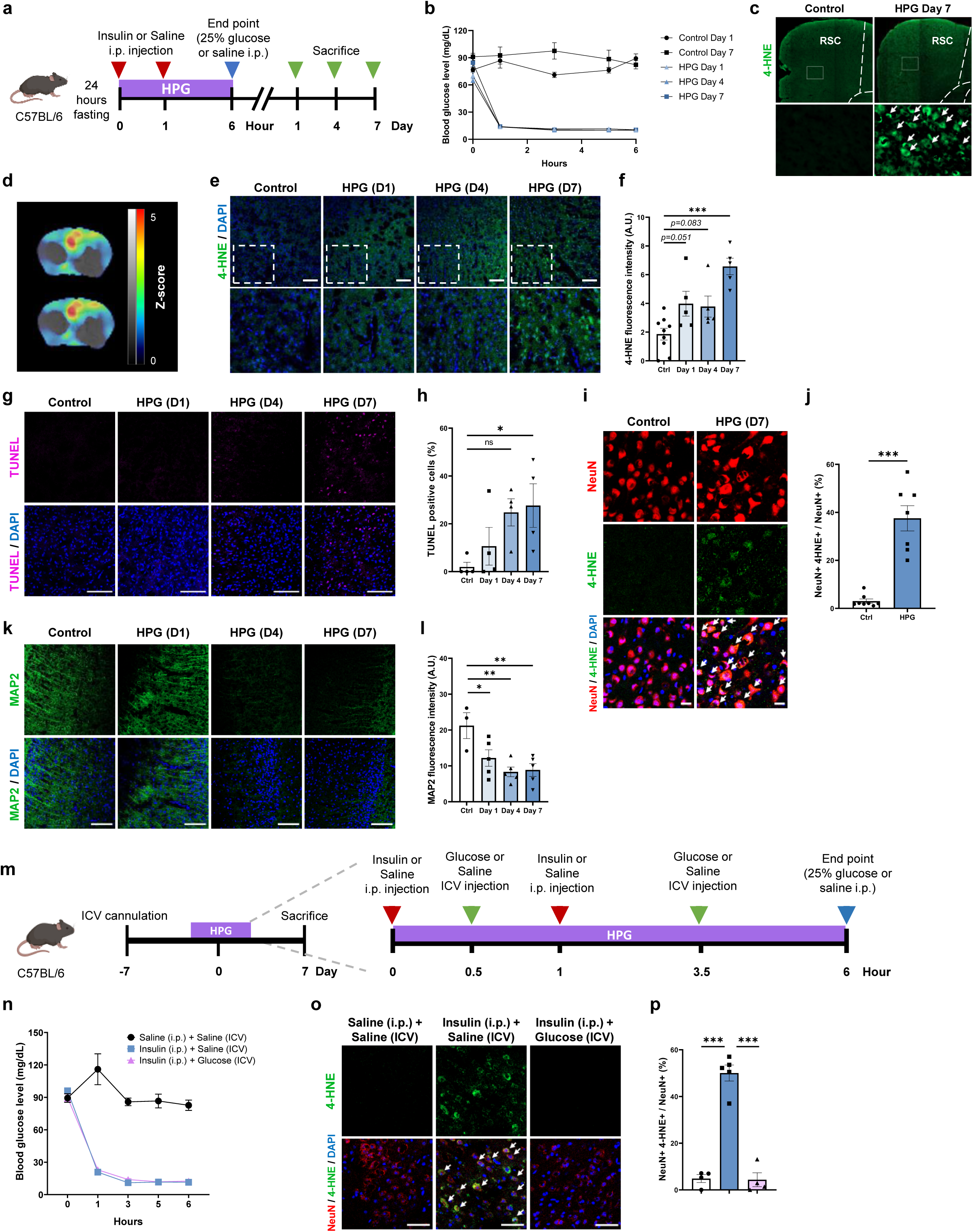
Oxidative stress, apoptosis, and neuronal dendritic loss in the RSC develop progressively after acute hypoglycemia. (a) Experimental design for hypoglycemia studies. (b) Blood glucose levels. (c) Unbiased screening of the cortex region with 4-HNE staining on day 7. White arrows indicate 4-HNE^+^ in the RSC. (d) Statistical parametric map of [^18^F]-FDG PET/CT image illustrating brain regions with significantly higher glucose uptake in the control group compared with the hypoglycemia group (*n* = 5 mice per group, p < 0.01). (e) Representative images of the RSC region with 4-HNE staining. Scale bars, 100 μm. (f) Quantification of 4-HNE intensity (*n* = 5 or 9 mice per group). (g) Representative images of the RSC region with TUNEL staining. Scale bars, 100 μm. (h) Quantification of TUNEL^+^ cells as a percentage of DAPI^+^ cells in the RSC region (*n* = 4 mice per group). (i) Representative images of NeuN and 4-HNE staining. Scale bars, 20 μm. White arrows indicate 4-HNE^+^ cells co-localized with NeuN^+^ cells. (j) Quantification of the percentage of 4-HNE^+^ cells co-localized with NeuN^+^ cells (*n* = 7 or 8 mice per group). (k) Representative images of MAP2 staining. Scale bars, 100 μm. (l) Quantification of MAP2 intensity in the RSC region (n = 3 or 5 mice per group). (m) Experimental design for glucose intracerebroventricular (ICV) supplementation. (n) Blood glucose levels. (o) Representative images of NeuN and 4-HNE staining. Scale bars, 50 μm. White arrows indicate 4-HNE^+^ cells co-localized with NeuN^+^ cells. (p) Quantification of the percentage of 4-HNE^+^ cells co-localized with NeuN^+^ cells (*n* = 4 or 5 mice per group). Data are presented as means ± SEM and were analyzed by (f, h, l, and p) one-way ANOVA followed by Dunnett’s multiple comparisons test, and (j) two-tailed Student’s *t* test. *P < 0.05; **P < 0.01; ***P < 0.001; ns, not significant.

We subsequently conducted time series analyses of the RSC following acute hypoglycemia on days 1, 4 and 7. We observed a gradual increase in the 4-HNE signal in the RSC, which reached significance on day 7 (Fig. 1e,f). To evaluate apoptosis, TdT-mediated dUTP nick end labeling (TUNEL) staining of the RSC was performed. The number of TUNEL^+^ cells progressively increased, with a significant change observed only on day 7 (Fig. 1g,h). These results indicate that oxidative damage and apoptosis in the RSC develop progressively after acute hypoglycemia.

To identify the specific cell types in which oxidative damage occurs due to hypoglycemia, we assessed the specific cell type in which 4-HNE staining increased in the RSC using markers for neurons (neuronal nuclei antigen, NeuN), astrocytes (glial fibrillary acidic protein, GFAP), and microglia (ionized calcium-binding adapter molecule 1, Iba-1). Notably, the ratio of 4-HNE^+^ cells to neurons significantly increased in the RSC at 7 days after severe hypoglycemia (Fig. 1i,j), whereas an increase in 4-HNE staining was not observed in cells positive for either GFAP or Iba-1 (Supplementary Fig. 1j-m). These findings reveal that oxidative damage in the RSC due to severe hypoglycemia occurs mainly in neurons rather than in astrocytes or microglia.

To further investigate neuronal loss in the RSC, we conducted a comprehensive analysis of markers indicating the neuronal soma, dendrites, and synapses. The fluorescence intensity of microtubule-associated protein 2 (MAP2; a cytoskeletal element) was significantly decreased on day 1 and was further decreased on day 7 (Fig. 1k,l). The intensity of the NeuN signal gradually decreased, although a significant change was not observed (Supplementary Fig. 1n,o). The expression levels of the synaptic proteins postsynaptic density protein-95 (PSD95) and synaptophysin were not significantly decreased during the progression of damage after acute hypoglycemia (Supplementary Fig. 1p,q). These results indicate that dendrites may be more susceptible than soma to acute hypoglycemia-induced damage in the RSC.

To determine whether hypoglycemia-induced brain damage is caused by glucose depletion in the brain or by other peripheral factors, we intracerebroventricularly injected glucose during severe hypoglycemia to maintain glucose levels in the brain without affecting peripheral blood glucose levels (Fig. 1m,n). Intracerebroventricular glucose injection significantly reduced the ratio of 4-HNE^+^ cells to neurons in the RSC after severe hypoglycemia compared to vehicle injection (Fig. 1o,p). These results demonstrate that brain damage after severe hypoglycemia is related to glucose deprivation in the brain *per se*, which can be prevented by timely supplementation of glucose in the brain.

### Mitochondrial fission, microglial activity, and IL-1β levels are increased in the RSC in the early stages of hypoglycemia-induced brain damage

Having established that neurons in the RSC are particularly vulnerable to acute hypoglycemia-induced damage, we next investigated the underlying cellular mechanisms. To this end, we performed single nucleus RNA sequencing (snRNA-seq) from the RSC of control and hypoglycemia-experienced mice on day 1. After quality control procedures (Supplementary Fig. 2a-d), cells were embedded, clustered, and annotated by canonical markers (Fig. 2a,b). Cell proportions of neuronal clusters decreased in the hypoglycemia group (Fig. 2c). These results suggest that hypoglycemia altered cell-type composition due to damage progression. Analysis of differentially expressed genes (DEGs) revealed significant upregulation of mitochondria-related genes in hypoglycemia-induced RSC (Fig. 2d). Gene ontology (GO) enrichment analysis of pseudo-bulk transcriptomes showed that upregulated genes were enriched for mitochondria-related biological processes, including oxidative phosphorylation and proton motive force-driven mitochondrial ATP synthesis (Fig. 2e, Supplementary Fig. 2e). Mitochondria-related biological processes were enriched predominantly in neurons and increased after hypoglycemia (Fig. 2f). These findings indicate that mitochondrial perturbation occurs primarily in neurons as an early pathological feature of hypoglycemic injury.

**Fig. 2.**
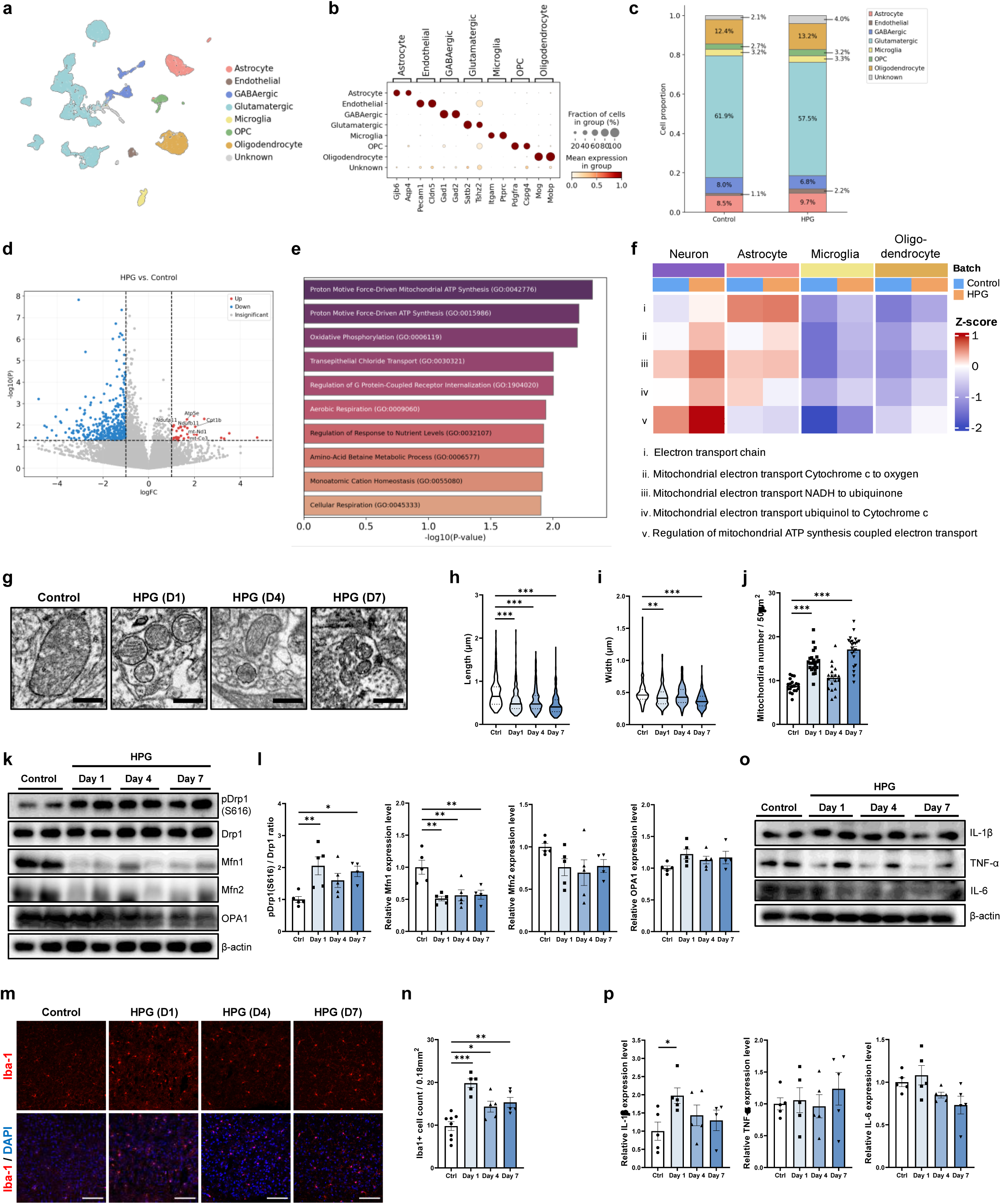
Mitochondrial fission, microglial activity, and IL-1β levels are increased in the RSC in the early stages of hypoglycemia-induced brain damage. (a) Uniform Manifold Approximation and Projection (UMAP) visualization of the snRNA-seq dataset (9,470 cells from 2 groups, *n* = 4 mice per group), colored by cell type. (b) Dot plot showing the expression of representative marker genes for each cell type. (c) Cell type proportions. (d) Volcano plot of DEGs. (e) GO enrichment analysis of genes upregulated in the HPG sample. (f) Gene set scoring-based heatmap of mitochondria-related GO terms across cell types. (g) Representative TEM images in the RSC. Scale bars, 0.5 μm. (h and i) Mitochondrial length in longitudinal sections (h) and width in cross sections (i). Quantification of mitochondria (*n* = 165-294) from 2 or 3 biologically independent samples per group. Data are presented as the median (solid line) with quartiles (dotted line) and whiskers showing minimum to maximum in the violin plots. (j) The number of mitochondria was counted in an area size of 50 μm^2^ (*n* = 18-24 independent images). Each dot represents the quantification of a single TEM image from 2 or 3 biologically independent samples in each group. (k and l) Representative immunoblots (k) and quantification (l) of mitochondrial dynamics proteins from the RSC tissues. (*n* = 4 or 5 mice per group). (m) Representative images of the RSC region with Iba-1 staining. Scale bars, 100 μm. (n) Quantification of Iba-1^+^ cells normalized by the area size of 0.18 mm^2^ (*n* = 5 or 8 mice per group). (o and p) Representative immunoblots (o) and quantification (p) of cytokine proteins from the RSC (*n* = 4 or 5 mice per group). Data are presented as means ± SEM and were analyzed by one-way ANOVA followed by Dunnett’s multiple comparisons test. *P < 0.05; **P < 0.01; ***P < 0.001.

To further assess mitochondrial abnormalities, we examined mitochondrial ultrastructure in the RSC by transmission electron microscopy (TEM) and observed morphological alterations (Fig. 2g). Significant decreases in both the length and width of the mitochondria were observed in the hypoglycemia group compared with the control group on days 1 and 7 (Fig. 2h,i). The number of mitochondria was also significantly increased on days 1 and 7 (Fig. 2j). To further assess alterations in mitochondrial dynamics in the RSC after hypoglycemia, we analyzed the protein expression of molecular markers of mitochondrial fission (dynamin-related protein 1, Drp1; activated form, pDrp1) and fusion (mitofusin1, Mfn1; mitofusin2, Mfn2; optic atrophy-1, OPA1). The pDrp1(S616)/Drp1 ratio in the RSC was significantly increased on day 1, indicating increased mitochondrial fission^16^, and this change remained significant on day 7 (Fig. 2k,l). However, the expression level of Mfn1 was significantly decreased on days 1, 4, and 7, whereas the expression levels of Mfn2 and OPA1 did not significantly change (Fig. 2k,l). These results indicate that acute hypoglycemia promotes mitochondrial fission and suppresses mitochondrial fusion in the RSC in the early stages of hypoglycemia-induced brain damage.

We also observed upregulation of genes involved in cellular stress response and proteostasis pathways, including protein ubiquitination and ubiquitin-dependent protein catabolic processes in the microglia cluster from snRNA-seq analysis (Supplementary Fig. 2f). To further examine inflammatory activity, we quantified microglial marker and cytokine levels in the RSC. The number of Iba-1^+^ cells was significantly increased in the RSC on day 1, as well as on days 4 and 7 (Fig. 2m,n). Since activated microglia are known to communicate with neurons via cytokines^17^, we next investigated the levels of cytokines that may mediate hypoglycemia-induced neuronal damage in the presence of reactive microglia. Notably, the expression level of IL-1β was significantly doubled on day 1 in the RSC, whereas the levels of TNF-α and IL-6 were not significantly altered during the progression of hypoglycemia-induced damage (Fig. 2o,p). These results implicate enhanced protein quality control and stress adaptation mechanisms in microglia, suggesting that microglia actively respond to severe hypoglycemia. These findings demonstrate that increased mitochondrial fission, inflammatory activation, and IL-1β expression precede brain damage in the RSC after acute hypoglycemia.

To ascertain whether increased mitochondrial fission and inflammatory activity precede brain damage, we examined the RSC of mice subjected to severe hypoglycemia for a relatively short period (3 hours of blood glucose levels below 20 mg/dL) (Supplementary Fig. 2g,h). We observed no significant difference between the groups in either the expression of an oxidative stress marker (4-HNE) or the expression of a dendritic marker (MAP2) in the RSC (Supplementary Fig. 2i-l). However, notably, the number of Iba-1^+^ microglia was significantly increased in the RSC in the hypoglycemia group (Supplementary Fig. 2m,n). We also observed elevated mitochondrial fission (increased pDrp1(S616)/Drp1 ratio) (Supplementary Fig. 2o,p). These results confirm that microglial activation and mitochondrial fission are increased before the development of significant brain damage in the RSC after acute hypoglycemia.

### Crosstalk between neuronal mitochondrial fission and microglial activation triggers hypoglycemia-induced neuronal damage in the RSC

Since both mitochondrial fission and inflammatory activation are increased in the RSC in the early stages of hypoglycemia-induced brain damage, we next investigated the causal relationship between mitochondrial fission and inflammatory activation. To this end, we treated mice with a mitochondrial division inhibitor (mdivi-1), a mitochondrial fission inhibitor, or minocycline, a tetracycline antibiotic that ameliorates microglial activity, during and after hypoglycemia (Fig. 3a). Neither mdivi-1 nor minocycline affected blood glucose levels during the period of hypoglycemia (Fig. 3b). The ratio of 4-HNE^+^ cells to neurons was significantly increased after hypoglycemia, and this change could be rescued by treatment with mdivi-1 or minocycline (Fig. 3c,d). Notably, the number of Iba-1^+^ microglia was decreased not only by minocycline but also by mdivi-1 (Fig. 3e,f). We further analyzed the morphology of microglia to evaluate microglial activation. Significant decreases in the number of branches and path length were observed in the vehicle-treated hypoglycemia group compared with the control group, suggesting microglial reactivity^18^, whereas the administration of either minocycline or even mdivi-1 caused the morphology of microglia to be comparable to that in the control group (Fig. 3g-i). Considering the increase in mitochondrial fission in the RSC after hypoglycemia (Fig. 2), we further identified the specific cell type in which mitochondrial fission occurred. An increase in the number of pDrp1(S616)^+^ cells was observed in the vehicle-treated hypoglycemia group, and interestingly, its pDrp1(S616) was significantly colocalized with NeuN (Fig. 3j,k) but not with GFAP (Supplementary Fig. 3a,b) or Iba-1 (Supplementary Fig. 3c,d). The ratio of pDrp1(S616)^+^ cells to neurons was significantly decreased in the mdivi-1-treated group and even in the minocycline-treated group (Fig. 3j,k). These results reveal that hypoglycemia-induced mitochondrial fission predominantly occurs in neurons rather than in astrocytes or microglia and that blocking this effect may prevent hypoglycemia-induced neuronal damage. Inhibition of either microglial activation or neuronal mitochondrial fission attenuated the other and rescue neuronal damage, implicating crosstalk between neuronal mitochondrial fission and microglial activation after severe hypoglycemia.

**Fig. 3.**
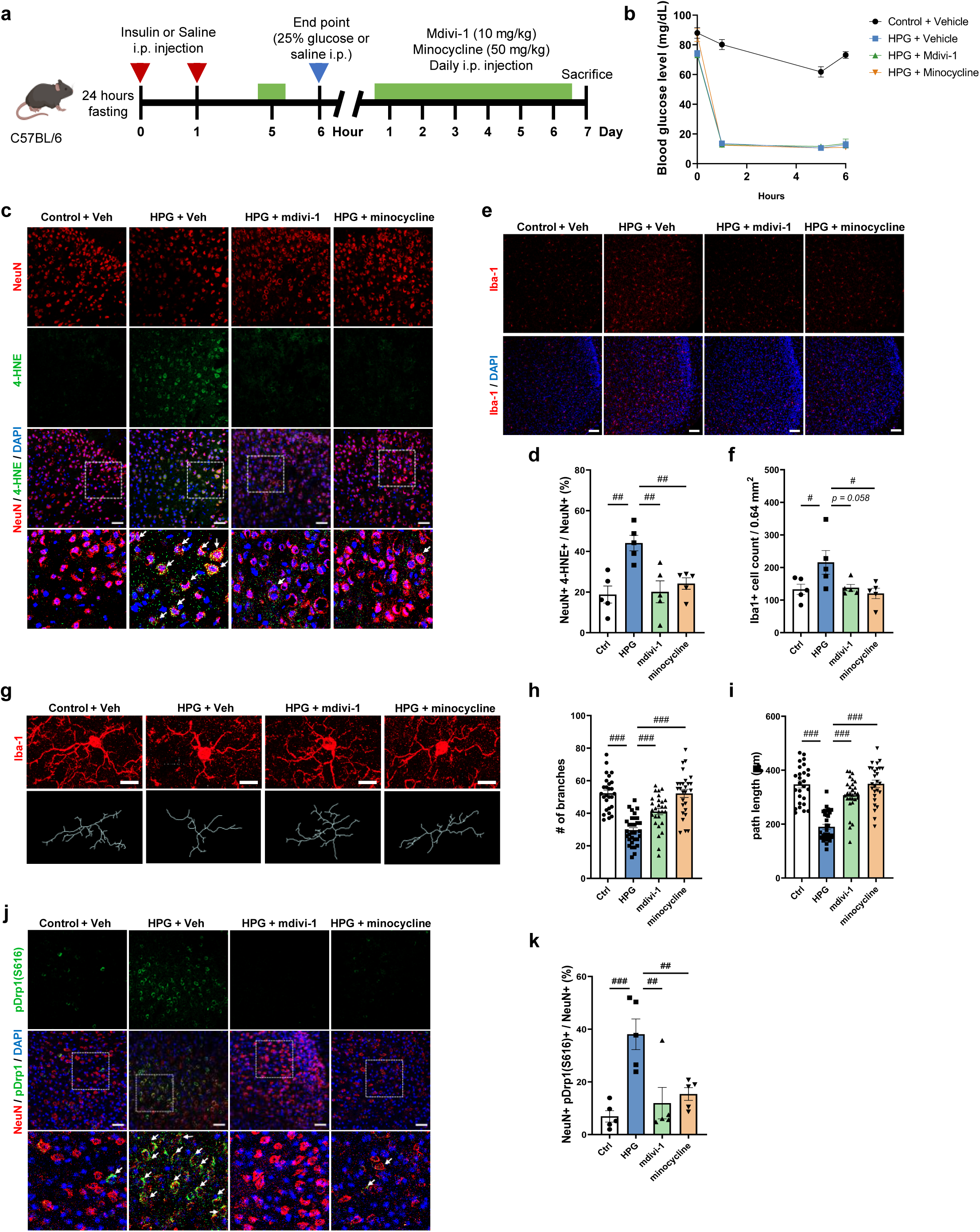
Crosstalk between neuronal mitochondrial fission and microglial activation triggers hypoglycemia-induced neuronal damage in the RSC. (a) Experimental design for mdivi-1 and minocycline treatment. (b) Blood glucose levels. (c) Representative images of NeuN and 4-HNE co-staining in the RSC. Scale bars, 50 μm. White arrows indicate 4-HNE^+^ cells co-localized with NeuN^+^ cells. (d) Quantification of the percentage of 4-HNE^+^ cells co-localized with NeuN^+^ cells (*n* = 5 mice per group). (e) Representative images of Iba-1 staining in the RSC. Scale bars, 100 μm. (f) Quantification of Iba-1^+^ cells normalized by the area size of 0.64 mm^2^ (*n* = 5 mice per group). (g-i) Representative Z-stack images of Iba-1^+^ microglia in the RSC (g). Scale bars, 10 μm. The branched paths of microglia were selected based on the captured images. Quantification of microglial branch number (h) and path length (i) (*n* = 27-32 independent microglia). Each dot represents the quantification of a single microglial cell from *n* = 5 biologically independent samples in each group. (j) Representative images in the RSC sections stained with NeuN and pDrp1(S616). White arrows indicate pDrp1^+^ cells co-localized with NeuN^+^ cells. Scale bars, 50 μm. (k) Quantification of the percentage of pDrp1^+^ cells co-localized with NeuN^+^ cells (*n* = 5 mice per group). Data are presented as means ± SEM and were analyzed by one-way ANOVA followed by Dunnett’s multiple comparisons test. #P < 0.05; ##P < 0.01; ###P < 0.001.

To examine whether the same mechanisms occur in female mice, we assessed hypoglycemia-induced neuronal damage in females. The ratios of 4-HNE^+^ and pDrp1(S616)^+^ cells to neurons were significantly increased in the RSC after hypoglycemia (Supplementary Fig. 3e-h). In addition, the number of Iba-1^+^ microglia was significantly increased in the hypoglycemia group (Supplementary Fig. 3i,j). These findings indicate that neuronal mitochondrial fission and microglial activation accompany hypoglycemia-induced damage in the female RSC, consistent with our observation in male mice.

We next tested whether our findings are preserved in a disease-relevant context by employing a streptozotocin (STZ)-induced diabetic mouse model, which mimics insulin-treated diabetes. Mice were injected with STZ and maintained under diabetes conditions for 10 weeks prior to the hypoglycemia experiment (Supplementary Fig. 3k). The ratio of 4-HNE^+^ cells to neurons was not significantly increased in the diabetes control group compared to the non-diabetes control group, indicating that the diabetic condition *per se* did not induce substantial oxidative stress in the RSC (Supplementary Fig. 3l,m). However, hypoglycemia in diabetic mice significantly increased the ratio of 4-HNE^+^ cells to neurons compared with the diabetes control group (Supplementary Fig. 3l,m). Similarly, the ratio of pDrp1(S616)^+^ cells to neurons and the number of Iba-1^+^ microglial cells were significantly increased in the hypoglycemia-experienced diabetic group. (Supplementary Fig. 3n-q). To validate the therapeutic effect of targeting mitochondrial fission in this model, we administered mdivi-1 or vehicle in the hypoglycemia-experienced diabetic group (Supplementary Fig. 3r). The ratios of 4-HNE^+^ cells and pDrp1(S616)^+^ cells to neurons were significantly decreased in the mdivi-1-treated hypoglycemia group compared to the vehicle-treated group (Supplementary Fig. 3s-v). The number of Iba-1^+^ microglial cells was also significantly decreased in the mdivi-1-treated hypoglycemia group (Supplementary Fig. 3w,x). These results demonstrate that the neuronal and microglial responses observed after acute hypoglycemia in normoglycemic mice are preserved under diabetic conditions, and that pharmacological inhibition of mitochondrial fission effectively attenuates hypoglycemia-induced brain damage even in a diabetes model.

### *In vitro* studies confirm cell type-specific mechanisms of hypoglycemia-induced neuronal damage via crosstalk with microglia

To further validate the cell type-specific mechanisms by which targeting mitochondrial dynamics or inflammation protects against hypoglycemia-induced neuronal damage, we used an *in vitro* system consisting of defined cell populations. In this model, cytotoxicity significantly increased when SH-SY5Y neuronal cells were exposed to glucose deprivation for 24 hours (Supplementary Fig. 4a). However, this effect was significantly ameliorated when mdivi-1 was added to the glucose-deprived medium (Supplementary Fig. 4b). We next employed a Transwell co-culture system of SH-SY5Y cells and BV-2 microglia to assess cell type-specific regulatory effects using mdivi-1 or minocycline (Fig. 4a). Pretreating SH-SY5Y cells directly with mdivi-1 significantly reduced apoptosis, as measured by the number of cleaved caspase-3^+^ (cCasp3^+^) SH-SY5Y cells, whereas minocycline pretreatment had no effect on SH-SY5Y cells (Fig. 4b,c). The decrease in mitochondrial length caused by glucose deprivation, which was detected by MitoTracker staining, was significantly prevented when SH-SY5Y cells were pretreated with mdivi-1 but not with minocycline (Fig. 4d,e). However, pretreating BV-2 cells with mdivi-1 did not prevent the increase in the number of cCasp3^+^ SH-SY5Y cells, suggesting that the effect of mdivi-1 was not mediated through microglia (Supplementary Fig. 4c-e). To exclude the possibility that minocycline directly regulates mitochondrial fission in neurons, minocycline was added to the glucose-deprived medium of SH-SY5Y cells not co-cultured with BV-2 cells (Supplementary Fig. 4f). Minocycline did not prevent the increase in the number of cCasp3^+^ SH-SY5Y cells or excessive mitochondrial fission (Supplementary Fig. 4g-j). These results indicate that mitochondrial fission in neurons, but not in microglia, is the key downstream regulator of neuronal damage following hypoglycemia. Furthermore, minocycline treatment reduced neuronal damage by inhibiting microglial activation but not by directly regulating mitochondrial fission in neurons in the hypoglycemia group.

**Fig. 4.**
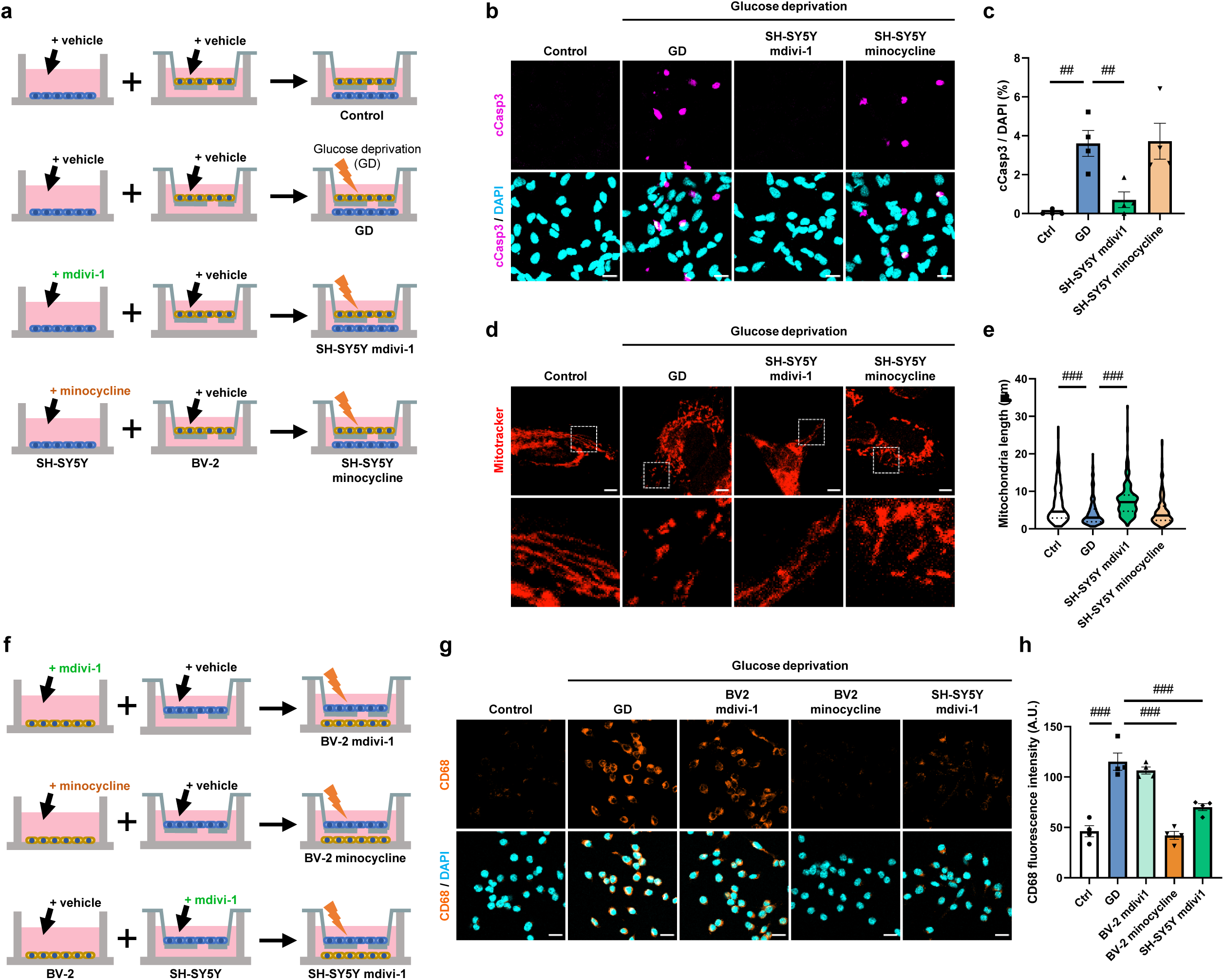
In vitro studies confirm cell type-specific mechanisms of hypoglycemia-induced neuronal damage via crosstalk with microglia. (a) Experimental design for mdivi-1 or minocycline pretreatment in SH-SY5Y cells. (b) Representative images of SH-SY5Y cells with cleaved caspase-3 staining. Scale bars, 20 μm. (c) Quantification of cleaved caspase-3^+^ cells as a percentage of DAPI^+^ cells (*n* = 4 independent samples per group). (d) Representative images of SH-SY5Y cells with MitoTracker. Scale bars, 5 μm. (e) Quantification of mitochondrial length (*n* = 165-321 mitochondria from 4 independent samples per group). Data are presented as the median (solid line) with quartiles (dotted line) and whiskers showing minimum to maximum in the violin plots. (f) Experimental design for mdivi-1 or minocycline pretreatment in BV-2 cells. (g) Representative images of BV-2 cells with CD68 staining. Scale bars, 20 μm. (h) Quantification of CD68 intensity (*n* = 4 independent samples per group). Data are presented as means ± SEM (c and h) and were analyzed by one-way ANOVA followed by Dunnett’s multiple comparisons test (c, e, and h). ##P < 0.01; ###P < 0.001.

Mitochondrial fission was increased mainly in neurons after hypoglycemia, whereas the extent of mitochondrial fission in microglia was comparable between the hypoglycemia group and the control group (Fig. 3). To further investigate whether mdivi-1 directly prevents microglial activation, BV-2 cells were treated with either mdivi-1 or minocycline during glucose deprivation (Supplementary Fig. 4k). The mean fluorescence intensity of CD68, a marker of activated microglia, was significantly lower in minocycline-treated BV-2 cells than in vehicle-treated cells after glucose deprivation, whereas mdivi-1 treatment had no effect (Supplementary Fig. 4l,m). These results imply that mdivi-1 does not directly suppress microglial activation. To further investigate whether the anti-inflammatory effects of mdivi-1 *in vivo* (Fig. 3e-i) involve neuron-to-microglia crosstalk, we examined whether mdivi-1 treatment targeting neurons indirectly modulates microglial activation. To this end, we once again employed a Transwell co-culture system to study the cell type-specific effects of mdivi-1 and minocycline (Fig. 4f). Pretreating BV-2 cells with minocycline significantly prevented the glucose deprivation-induced increase in the CD68 intensity, whereas pretreating BV-2 cells with mdivi-1 did not (Fig. 4g,h). Notably, pretreating SH-SY5Y cells with mdivi-1 significantly reduced the CD68 intensity (Fig. 4g,h), suggesting that there is crosstalk between neuronal mitochondrial fission and microglial activation. Therefore, mdivi-1 treatment prevented microglial activation through its interaction with the inhibition of neuronal mitochondrial fission but not through direct regulation of microglial activation in the hypoglycemia group. These *in vitro* findings confirm that the effects of mdivi-1 and minocycline observed *in vivo* (Fig. 3) are attributable to their cell type-specific, on-target mechanisms.

### IL-1 signaling mediates the crosstalk of microglial activation with neuronal mitochondrial fission to trigger hypoglycemia-induced neuronal damage in the RSC

As we observed an elevated level of IL-1β in the RSC (Fig. 2), we hypothesized that activated microglia release IL-1β, which may exacerbate mitochondrial fission in neurons. To verify this hypothesis, we administered intracerebroventricular injections of interleukin-1 receptor antagonist (IL-1ra) during and after severe hypoglycemia. While this treatment did not affect blood glucose levels (Fig. 5a,b), the ratio of 4-HNE^+^ cells to neurons was significantly decreased in the IL-1ra-injected hypoglycemia group compared with the vehicle-injected hypoglycemia group (Fig. 5c,d). Notably, the ratio of pDrp1(S616)^+^ cells to neurons (Fig. 5e,f) and the number of Iba-1^+^ microglial cells (Fig. 5g,h) were also significantly decreased in the IL-1ra-injected hypoglycemia group. Moreover, microglial morphology analysis revealed that the reduction in the number of branches and decrease in the process length of activated microglia after hypoglycemia were significantly rescued by IL-1ra injection (Fig. 5i-k). These results imply that activation of the IL-1 signaling pathway by acute hypoglycemia may mediate neuronal damage by regulating neuronal mitochondrial fission and microglial activity.

**Fig. 5.**
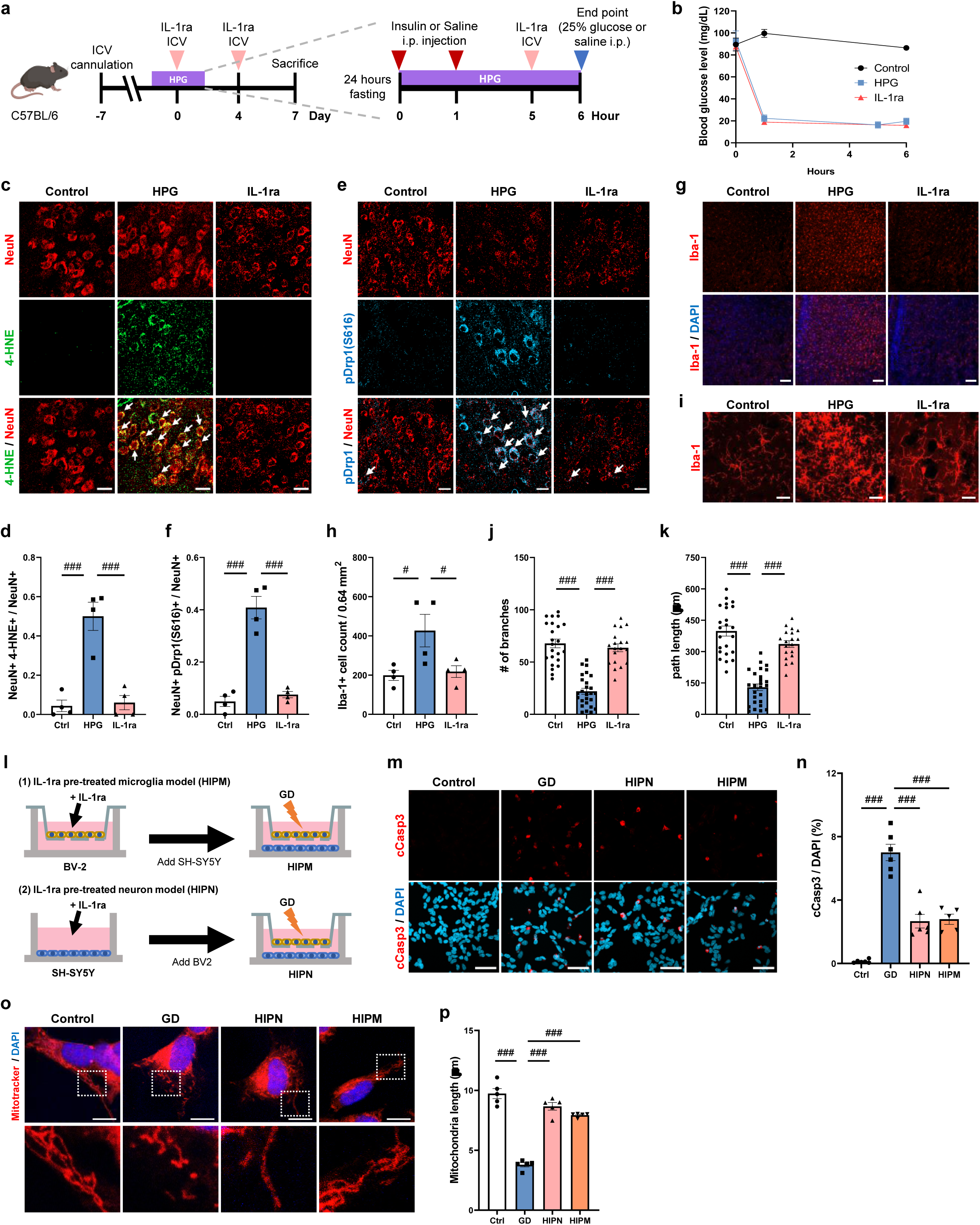
IL-1 signaling mediates the crosstalk of microglial activation with neuronal mitochondrial fission to trigger hypoglycemia-induced neuronal damage in the RSC. (a) Experimental design for IL-1ra ICV injection. (b) Blood glucose levels. (c) Representative images of the RSC tissues with NeuN and 4-HNE co-staining. White arrows indicate 4-HNE^+^ cells co-localized with NeuN^+^ cells. Scale bars, 20 μm. (d) Quantification of the percentage of 4-HNE^+^ cells co-localized with NeuN^+^ cells (*n* = 4 mice per group). (e) Representative images of the RSC tissues with pDrp1(S616) and NeuN co-staining. White arrows indicate pDrp1^+^ cells co-localized with NeuN^+^ cells. Scale bars, 20 μm. (f) Quantification of the percentage of pDrp1^+^ cells co-localized with NeuN^+^ cells (*n* = 4 mice per group). (g) Representative images of Iba-1 staining. Scale bars, 100 μm. (h) Quantification of Iba-1^+^ cells normalized by the area of 0.64 mm^2^ (*n* = 4 mice per group). (i) Representative Z-stack images in the RSC sections with Iba-1 staining. Scale bars, 20 μm. (j and k) Quantification of the number of branches (j) and path length (k) (*n* = 19-26 independent microglia). Each dot represents the quantification of a single microglial cell from *n* = 4 biologically independent samples in each group. (l) Experimental design for IL-1ra pretreatment in BV-2 or SH-SY5Y cells. (m) Representative images of SH-SY5Y cells with cleaved caspase-3 staining. Scale bars, 50 μm. (n) Quantification of cleaved caspase-3^+^ cells as a percentage of DAPI^+^ cells (*n* = 5 or 6 independent samples per group). (o) Representative Z-stack images of SH-SY5Y cells with MitoTracker staining. Scale bars, 10 μm. (p) Quantification of mitochondrial length (*n* = 5 independent samples per group). Data are presented as means ± SEM and were analyzed by one-way ANOVA followed by Dunnett’s multiple comparisons test. #P < 0.05; ###P < 0.001.

We further investigated whether these effects of blocking IL-1 signaling involve the direct regulation of mitochondrial fission in neurons or the indirect regulation of mitochondrial fission through the inhibition of microglial activation. To explore this, we again utilized a Transwell co-culture system and modulated IL-1 signaling in specific cell types. We pretreated SH-SY5Y or BV-2 cells with IL-1ra and then co-cultured them in a Transwell system with glucose-deprived medium (Fig. 5l). The number of cCasp3^+^ SH-SY5Y cells was significantly reduced when IL-1ra was applied to either SH-SY5Y or BV-2 cells (Fig. 5m,n). To determine whether blocking IL-1 signaling prevents neuronal damage by regulating mitochondrial fission in neurons, we further analyzed alterations in mitochondrial dynamics through MitoTracker staining. The inhibition of IL-1 signaling specifically in either SH-SY5Y or BV-2 cells significantly prevented excessive mitochondrial fission in neurons (Fig. 5o,p). These results indicate that IL-1 signaling in both neurons and microglia plays a crucial role in modulating neuronal damage following hypoglycemia. Collectively, our data indicate that IL-1 signaling can both directly mediate mitochondrial fission in neurons and indirectly modulate mitochondrial fission through microglial activation after hypoglycemia, resulting in neuronal damage.

We next assessed whether blockade of IL-1 signaling could confer similar neuroprotective effects in hypoglycemia-experienced diabetic mice (Supplementary Fig. 5a). IL-1ra treatment significantly reduced the ratios of 4-HNE^+^ cells and pDrp1(S616)^+^ cells to neurons compared to the vehicle-treated group (Supplementary Fig. 5b-e). In parallel, the number of Iba-1^+^ microglial cells was also significantly decreased in the IL-1ra-treated hypoglycemia group (Supplementary Fig. 5f,g). These results suggest that pharmacological blockade of IL-1 signaling also mitigates hypoglycemia-induced brain damage in a diabetic model.

### The RSC-specific inhibition of mitochondrial fission and IL-1 signaling prevents hypoglycemia-induced damage in the RSC

Whereas the preceding *in vivo* experiments involved systemic or intracerebroventricular drug administration (Fig. 3a; Fig. 5a), we next implanted unilateral cannulas in the RSC to manipulate mitochondrial fission and IL-1 signaling in a region-specific manner (Supplementary Fig. 6a). Hypoglycemia-induced damage to the contralateral RSC was comparable among vehicle-, mdivi-1-, and IL-1ra-injected mice exposed to hypoglycemia, with no significant differences in the number of 4-HNE^+^ cells (Supplementary Fig. 6b,c). To evaluate the protective effects of mdivi-1 and IL-1ra, we calculated the ratio of the number of 4-HNE^+^ cells in the ipsilateral RSC to that in the contralateral RSC for each mouse. Cannulation and vehicle delivery *per se* resulted in a trend toward an increased number of 4-HNE^+^ cells (∼1.5-fold) in the ipsilateral compared to the contralateral side in both control and hypoglycemia groups, likely due to invasive cannula implantation (Supplementary Fig. 6d). This ratio was decreased by injection of either mdivi-1 or IL-1ra during and after hypoglycemia (Supplementary Fig. 6d). These data demonstrate that RSC-specific inhibition of mitochondrial fission or IL-1 signaling can prevent hypoglycemia-induced damage, consistent with the protective effects observed through systemic administration of these compounds.

### Drp1-dependent mitochondrial fission and IL-1 signaling engage in reciprocal regulation to drive hypoglycemia-induced neuronal damage in the RSC

To elucidate the mechanistic basis of the excessive mitochondrial fission observed in the RSC, we investigated the activity of Drp1, a key regulator of mitochondrial fission that initiates fission through its translocation to the outer mitochondrial membrane (Fig. 6a). To assess whether hypoglycemia triggers the pathological translocation of Drp1 to mitochondria, we isolated mitochondrial fractions from the RSC one day after the hypoglycemia experiment (Fig. 6b). Mitochondrial Drp1 levels were significantly increased in the hypoglycemia group compared to controls, confirming that Drp1 translocation to mitochondria underlies the fission observed after hypoglycemia (Fig. 6c,d). Importantly, mdivi-1 treatment suppressed this translocation, further supporting a Drp1-dependent mechanism for hypoglycemia-induced mitochondrial fission.

**Fig. 6.**
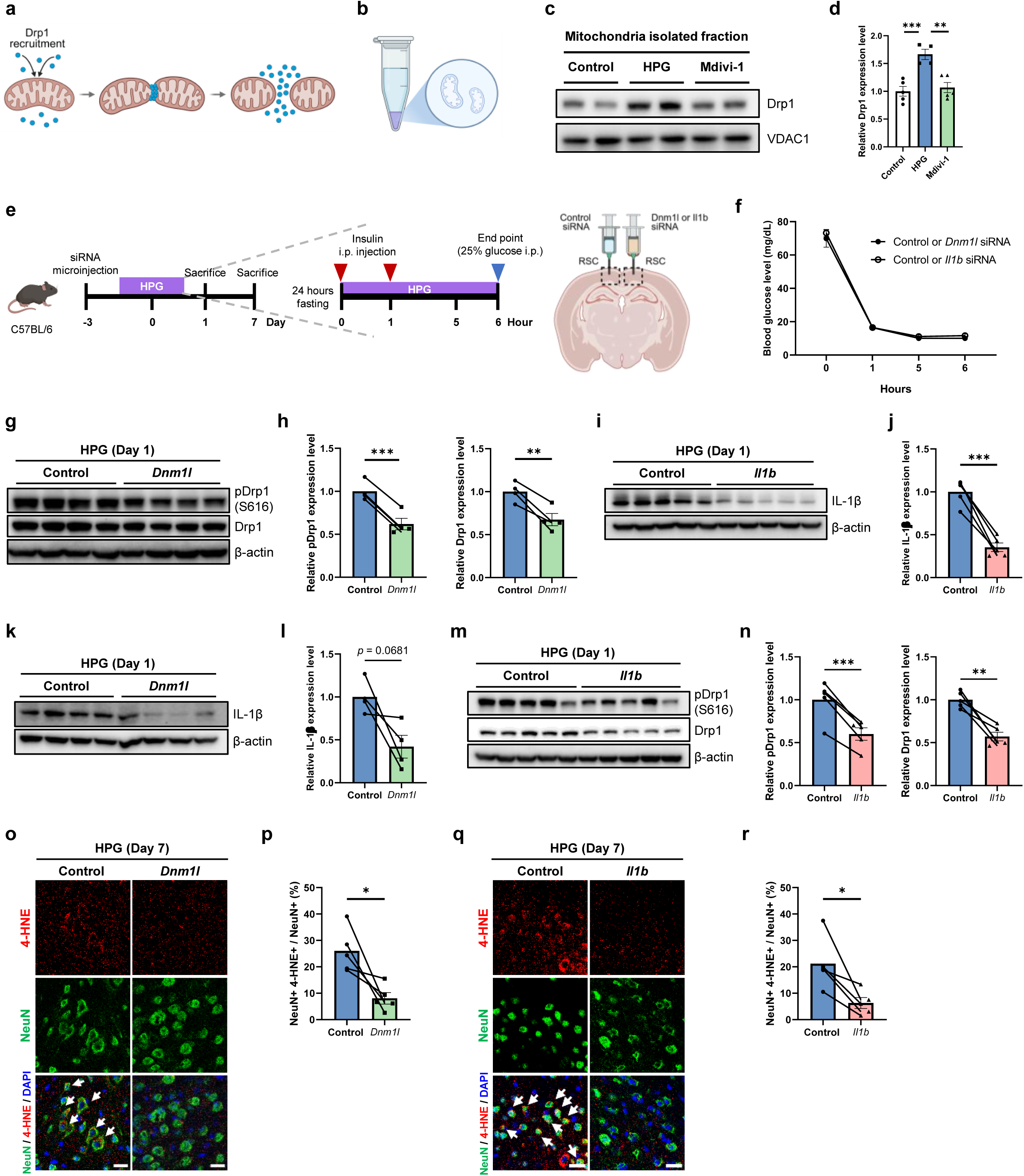
Drp1-dependent mitochondrial fission and IL-1 signaling engage in reciprocal regulation to drive hypoglycemia-induced neuronal damage in the RSC. (a) Schematic of Drp1-dependent mitochondrial fission. (b) Schematic of mitochondria isolation. (c and d) Immunoblot analysis from isolated mitochondria fraction of the RSC tissues (*n* = 4 or 5 mice). Data are presented as means ± SEM and were analyzed by one-way ANOVA followed by Tukey’s multiple comparisons test. **P < 0.01; ***P < 0.001. (e) Experimental design of the study. *Dnm1l* (*n* = 9 mice) or *Il1b* (*n* = 10 mice) siRNA was injected in the right RSC while control siRNA was injected in the contralateral RSC. (f) Blood glucose levels. (g-n) Immunoblot analysis from the RSC tissues with antibody against mitochondrial fission proteins (pDrp1(S616) and Drp1) and IL-1β on day 1 after hypoglycemia induction (*n* = 4 or 5 mice per group). (o-r) Immunohistochemistry analysis from the RSC tissues with NeuN and 4-HNE co-staining on day 7 after hypoglycemia induction (*n* = 5 mice per group). White arrows indicate 4-HNE^+^ cells co-localized with NeuN^+^ cells. Scale bars, 20 μm. Each dot represents a single RSC site from *n* = 4 or 5 mice injected unilaterally with *Dnm1l* or *Il1b* siRNA and contralaterally with control siRNA. Paired t-tests were used for statistical comparisons between hemispheres. Data are presented as means ± SEM. *P < 0.05; **P < 0.01; ***P < 0.001.

We next examined the functional relationship between Drp1-dependent mitochondrial fission and IL-1 signaling during hypoglycemia-induced neuronal damage in the RSC. To address this, we used *in vivo* siRNA-mediated knockdown of either *Dnm1l* (encoding Drp1) or *Il1b* (encoding IL-1β) within the RSC. In each mouse, target siRNA was injected into the right RSC and control siRNA into the contralateral left RSC, followed by the severe hypoglycemia experiment 3 days later (Fig. 6e,f). *Dnm1l* siRNA significantly reduced hypoglycemia-induced increases in pDrp1(S616) and Drp1 protein levels compared to the contralateral control siRNA-injected RSC at day 1 (Fig. 6g,h). Similarly, *Il1b* siRNA reduced IL-1β expression levels compared to the contralateral control side (Fig. 6i,j). Notably, knockdown of *Dnm1l* also suppressed IL-1β upregulation, while *Il1b* siRNA reduced pDrp1(S616) and Drp1 levels (Fig. 6k-n), suggesting that IL-1 signaling promotes mitochondrial fission in part by regulating Drp1. Together, these data support reciprocal crosstalk between Drp1-dependent mitochondrial fission and IL-1 signaling during hypoglycemia-induced neuronal damage. Finally, knockdown of either *Dnm1l* or *Il1b* significantly reduced the ratio of 4-HNE^+^ cells to neurons compared to the contralateral control siRNA-injected RSC (Fig. 6o-r). Collectively, these data support a feedforward mechanism in which hypoglycemia induces Drp1-dependent mitochondrial fission in neurons, thereby engaging microglial IL-1 signaling, which in turn further amplifies neuronal injury in the RSC.

### Regulation of excessive mitochondrial fission or IL-1 signaling prevents impairments in spatial learning and memory retrieval

The RSC is a critical region for spatial memory storage^19^ and retrieval^20^. Indeed, there have been reports indicating that severe hypoglycemia can clinically impact spatial memory in patients with diabetes^21,22^. Since we consistently observed damage in this region after hypoglycemia, we tested whether regulating mitochondrial fission in neurons is a possible therapeutic strategy to prevent the impairment of spatial learning and memory retrieval. To regulate mitochondrial fission directly or indirectly, we intraperitoneally injected mdivi-1 or IL-1ra, respectively (Fig. 7a). Spatial learning, memory, and memory retention were examined using the Morris water maze test (Fig. 7b). Compared with the control mice, the mice in the vehicle-treated hypoglycemia group presented a significantly longer escape latency during the training session, whereas mice in the mdivi-1-treated hypoglycemia group presented a significant decrease in escape latency (Fig. 7c). Compared with vehicle treatment, IL-1ra treatment also tended to decrease the escape latency. In the probe trial, the IL-1ra-treated hypoglycemia group spent significantly more time in the target quadrant than did the vehicle-treated hypoglycemia group. Compared with the vehicle-treated hypoglycemia group, both the IL-1ra- and mdivi-1-treated hypoglycemia groups presented a significant decrease in the mean distance required to reach the platform (Fig. 7d). In the memory retrieval trial, compared with the vehicle-treated hypoglycemia group, both treatment groups spent significantly more time in the target quadrant and exhibited a shorter mean distance required to reach the platform (Fig. 7e). These results suggest that hypoglycemia-induced impairment of spatial learning and memory can be prevented directly by regulating neuronal mitochondrial fission or indirectly by targeting IL-1 signaling.

**Fig. 7.**
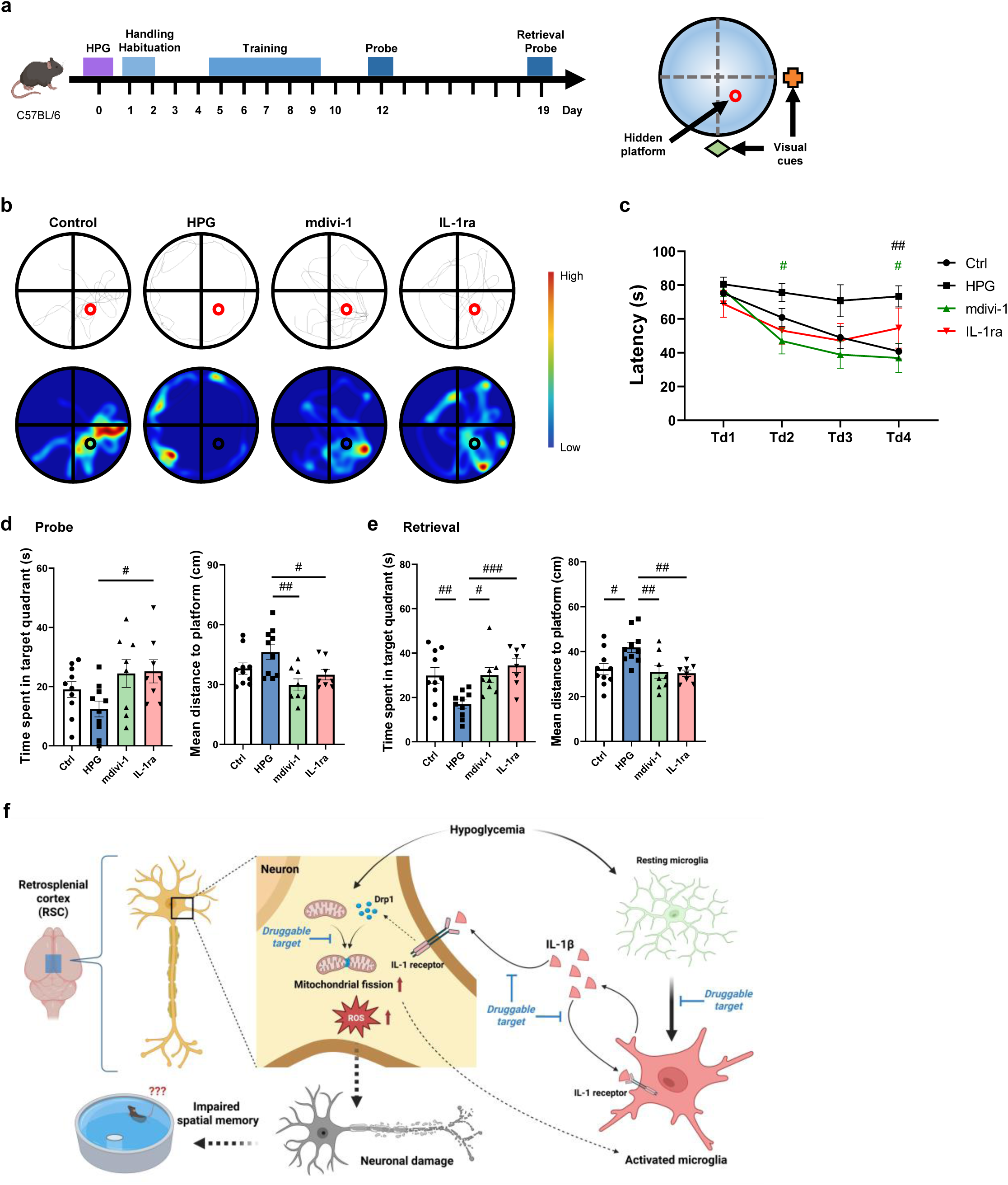
Regulation of excessive mitochondrial fission or IL-1 signaling prevents impairments in spatial learning and memory retrieval. (a) Experimental design of the Morris water maze and schematic representation of the maze. (b) Representative visualized track and heatmap during the probe trial. (c) Escape latency during the training session (*n* = 8 or 10 mice per group). (d and e) Quantification of the Morris water maze analysis in the probe trial (d) and the memory retrieval probe trial (e) (*n* = 8 or 10 mice per group). (f) Schematic representation showing how hypoglycemia-induced neuronal mitochondrial fission and microglial activation contribute to neuronal damage in the RSC and spatial memory impairment. Data are presented as means ± SEM and were analyzed by (c) two-way repeated-measures ANOVA followed by Dunnett’s multiple comparisons test and (d and e) one-way ANOVA followed by Dunnett’s multiple comparisons test. #P < 0.05; ##P < 0.01; ###P < 0.001.

The hippocampus is also known to be involved in spatial memory^23^, but we found that the hypoglycemia model mice and control mice exhibited a comparable level of oxidative damage in the hippocampus (Supplementary Fig. 1), suggesting that spatial memory deficits after hypoglycemia may not be attributed to hippocampal damage. Furthermore, in the open field test, elevated plus maze test, and tail suspension test, no significant alterations in behavior were observed in the vehicle-, mdivi-1-, or IL-1ra-treated hypoglycemia group compared with the control group (Supplementary Fig. 6e-h). These data indicate that the spatial memory impairment observed in the Morris water maze test may not have been affected by or caused by depression or anxiety.

## DISCUSSION

In this study, we report that acute severe hypoglycemia can selectively damage the RSC, among cortical and hippocampal regions, leading to spatial memory impairment (Fig. 7f). This brain damage develops progressively and is characterized by neuronal mitochondrial fission, microglial activation, and elevated IL-1β levels during the early stages. Notably, oxidative damage and mitochondrial fission occur predominantly in neurons rather than astrocytes or microglia in the RSC. Our findings reveal that bidirectional crosstalk between neuronal mitochondrial fission and microglial IL-1 signaling mediates this damage. Furthermore, hypoglycemia-induced neuronal damage and impaired cognitive function can be prevented either directly through the regulation of neuronal mitochondrial fission or indirectly through the inhibition of IL-1 signaling.

We identified the RSC as a previously unrecognized region vulnerable to severe hypoglycemia. The mechanisms underlying the particular vulnerability of this area to hypoglycemic stress remain an unresolved question. The RSC is well established as a crucial hub in various networks, including the default mode network, exhibiting extensive interconnectivity with various brain structures (e.g., the hippocampus, thalamus, midbrain, hypothalamus)^24,25^. Compared with other cortical areas, the ventral retrosplenial cortex has an exceptionally high vascular density^26^, suggesting elevated metabolic activity that requires greater glucose and oxygen consumption to support its function as a network hub. Indeed, studies have documented alterations in brain network activity and functional connectivity during hypoglycemia, particularly in patients with type 1 diabetes^27,28^. Even in patients with insulinoma, significant abnormalities in intra- and internetwork connectivity have been reported to be associated with cognitive impairment^29^. The vulnerability of the RSC to hypoglycemia-induced damage may be explained by the combined effects of extensive neural network disruption and inadequate metabolism during and after severe hypoglycemia, but the mechanism warrants further investigation.

Our findings demonstrate that hypoglycemia-induced mitochondrial fission occurs predominantly in neurons rather than in microglia or astrocytes, and that this fission is driven by Drp1-dependent mechanisms. This neuron-specific response may be attributed to the greater mitochondrial density in neurons than in glial cells^30^. Importantly, we show that the selective inhibition of Drp1-mediated mitochondrial fission in neurons is sufficient to prevent both neuronal damage and microglial activation. The mechanism underlying hypoglycemia-induced mitochondrial fission likely involves calcium dysregulation, as several studies have documented increased calcium influx in neurons during hypoglycemic conditions^31,32^. Given that elevated intracellular calcium levels are a trigger of mitochondrial fission^33^, our findings support a mechanistic model in which hypoglycemia-induced calcium influx drives mitochondrial fission in neurons, ultimately leading to neuronal injury and subsequent inflammation.

In this study, microglial modulation by minocycline treatment significantly attenuated hypoglycemia-induced mitochondrial fission in neurons. Minocycline, a tetracycline antibiotic, has been shown to inhibit microglial activation and reduce neuronal damage in the cortex and hippocampus in acute brain injury models^34–36^. However, its practical utility remains controversial, with evidence of effects ranging from neuroprotection to exacerbation of neurotoxicity in various experimental models and human trials^37,38^, possibly owing to its pleiotropic mode of action. This highlights the need to elucidate the precise mechanism by which microglial inhibition prevents hypoglycemia-induced neuronal damage. In this study, we also demonstrated that activated microglia regulate neuronal mitochondrial fission via IL-1 signaling, ultimately leading to neuronal damage. While the activation of IL-1 signaling is known to trigger neuronal damage^39^, inhibition of IL-1 signaling has been shown to exert protective effects in models of neurodegenerative diseases, traumatic brain injury, and ischemic brain damage^40–42^. Our findings show that both treatment of hypoglycemic mice with IL-1ra and neuron-specific inhibition of IL-1 signaling in glucose-deprived medium can prevent neuronal damage and mitochondrial fission. Importantly, genetic suppression of *Il1b* significantly reduced total Drp1 protein levels in the RSC following hypoglycemia, suggesting that IL-1 signaling promotes mitochondrial fission at least in part by upregulating Drp1 expression. Given that anakinra (recombinant IL-1ra) is already clinically approved and well tolerated in humans, repurposing this agent for neuroprotection in patients with diabetes who experience severe hypoglycemia represents a promising translational strategy. Nevertheless, the precise molecular mechanism by which the pathway downstream of IL-1 signaling regulates mitochondrial fission in neurons remains to be elucidated in future studies.

Our results reveal that neuron-specific regulation of mitochondrial fission prevents microglial activation after hypoglycemia, indicating a critical role for neuronal signals in modulating neuroinflammatory responses. This neuron-to-microglia communication aligns with previous findings across various neurological conditions, including neurodegenerative diseases, in which neurons regulate microglial activation through multiple molecular pathways^43,44^. Our findings expand upon this concept by demonstrating that inhibiting mitochondrial fission in neurons is sufficient to prevent microglial activation. Female mice exhibited the same RSC pathology as males following severe hypoglycemia, suggesting sex-independent mechanisms. This mechanistic insight could have important therapeutic implications, as it suggests that neuroprotective strategies targeting neuronal mitochondria might be sufficient to prevent both neuronal damage and neuroinflammation.

In our hypoglycemia mouse model, neuronal damage developed in a progressive manner, and the application of therapeutic interventions during this progressive period prevented hypoglycemia-induced brain damage. This suggests that a sufficient window exists for clinical intervention during recovery following severe hypoglycemic episodes. Our findings highlight that therapeutic interventions could prevent long-term neurological complications even after severe hypoglycemic events. Moreover, our mechanistic findings were recapitulated in STZ-induced diabetes mice, where inhibition of either mitochondrial fission or IL-1 signaling conferred neuroprotection, supporting the translational potential of targeting mitochondrial dynamics as a therapeutic strategy to prevent hypoglycemia-associated neurotoxicity in patients with diabetes treated with insulin.

Our findings also have broader implications for understanding how systemic metabolic perturbations can lead to selective neuronal vulnerability across brain regions. We highlight a bidirectional crosstalk between neuronal mitochondrial dynamics and microglial IL-1 signaling, both of which are perturbed in response to metabolic stress such as glucose deprivation. This reciprocal interaction may represent a fundamental mechanism by which metabolic abnormalities such as hypoglycemia, insulin resistance, or broader dysmetabolic states converge on the brain to drive neuroinflammation and neuronal injury. Given that mitochondrial dysfunction and immune activation are common features in diabetes, obesity, and other metabolic syndromes, the neuron– microglia interplay described here may underlie shared neuropathological outcomes across diverse conditions. Targeting this intercellular axis may offer a promising therapeutic strategy to mitigate brain complications associated with systemic metabolic dysregulation. Interestingly, RSC dysfunction has been implicated in various cognitive disorders, including Alzheimer’s disease^45^. While our study focused on metabolic stress-induced injury, future investigations may explore whether similar mitochondrial-microglial interactions contribute to RSC vulnerability in neurodegenerative contexts.

Our study has some limitations. To enable timely observation of hypoglycemia-induced damage mechanisms, we restricted our analyses to an acute hypoglycemia model. Whether accumulated mitochondrial fission and inflammatory activation contribute to progressive injury during recurrent hypoglycemic episodes remains unclear. Although we observed increased mitochondrial fission followed by oxidative damage in neurons after hypoglycemia, the direct causal relationship between mitochondrial fragmentation and oxidative damage requires further investigation. While systemic delivery of mitochondrial fission inhibitor and IL-1 receptor antagonist demonstrated neuroprotection in our study, potential effects on peripheral organs were not assessed. Moreover, human studies testing safety and pharmacokinetics should be performed to evaluate the potential clinical use.

In conclusion, our findings identify a feedforward mechanism underlying hypoglycemia-induced neuronal damage, in which Drp1-dependent mitochondrial fission in neurons and microglial IL-1 signaling reinforce one another to amplify neuronal injury. These findings suggest that targeting mitochondrial fission, either directly or indirectly through IL-1 signaling, may help prevent neuronal damage and spatial memory impairment after hypoglycemia.

## MATERIALS AND METHODS

### Animals

Six- to eight-week-old male and female C57BL/6 mice were purchased from Orient Bio (Gapyeong, Korea). All the mice were housed in a controlled environment on a 12-hour light/dark cycle at a constant temperature of 23 ± 1°C and 50 ± 10% relative humidity. The mice were given *ad libitum* access to a normal chow diet (Purina, Gyeonggi, Korea) and water. All experimental procedures were approved by the Institutional Animal Care and Use Committee of Seoul National University.

### Severe hypoglycemia induction

Nine-week-old male and female C57BL/6 mice were fasted overnight or for 24 hours. Hypoglycemia was induced by two intraperitoneal injections of 20 U/kg Humulin R (Eli Lilly and Company, U.S.) at a one-hour interval. Glucose levels in blood taken from the tail vein were monitored at 1- to 2-hour intervals using an ACCU-CHEK instant glucometer (Roche Diagnostics, Switzerland) to ensure sustained severe hypoglycemia (< 20 mg/dL glucose). To terminate hypoglycemia, 200 μl of 25% glucose was administered intraperitoneally at 6 hours after the first insulin injection. Under the same experimental conditions, euglycemic control mice were administered an equal volume of saline.

### Drug treatment

Mdivi-1 (Sigma‒Aldrich, M0199) and minocycline (Sigma‒Aldrich, M9511) were initially administered intraperitoneally 1 hour before the injection of 25% glucose and subsequently administered once daily for 6 days. Mdivi-1 was dissolved in dimethyl sulfoxide (DMSO) to prepare a stock solution, which was then diluted in sterile saline (4% final DMSO concentration) for injection. Owing to its low aqueous solubility, the solution was gently sonicated (30% amplitude, 30 sec), followed by immediate intraperitoneal injection. Minocycline was dissolved in sterile saline. The mice were administered mdivi-1 (10 mg/kg), minocycline (50 mg/kg), or vehicle.

For the Morris water maze test, a stock solution of IL-1ra (Cayman Chemical, 21349) was diluted in sterile saline (1% final ethanol concentration). The mice received intraperitoneal injections of mdivi-1 (10 mg/kg), IL-1ra (2 mg/kg), or vehicle for 7 consecutive days.

### Acute hypoglycemia experiment in STZ-induced diabetic mouse model

STZ was dissolved in 100 mM sodium citrate buffer (pH 4.5). Eight-week-old male C57BL/6 mice were i.p. injected with STZ (55 mg/kg) once daily for five consecutive days. Following diabetes induction (blood glucose > 200 mg/dL), the diabetic condition was maintained for 10 weeks prior to the hypoglycemia experiment. For the hypoglycemia experiment, mice were fasted for 24 hours and then intraperitoneally injected with 30 U/kg of Humulin R at one-hour intervals, followed by an additional 20 U/kg one hour later. Severe hypoglycemia (<20 mg/dL glucose) was maintained for 5 hours after the final insulin injection. Hypoglycemia was terminated by administration of 200 μl of 25% glucose solution. Mdivi-1 (10 mg/kg) or IL-1ra (2 mg/kg) were administered intraperitoneally 1 hour before the termination of hypoglycemia and subsequently once daily for 6 consecutive days. Control mice (non-diabetic or diabetic mice not subjected to insulin-induced hypoglycemia) received equivalent volumes of saline under the same experimental schedule.

### Stereotaxic surgery and intracerebroventricular injection

Eight-week-old male C57BL/6 mice were anesthetized, and stainless-steel guide cannulas (Protech International, C315GS, 3 mm projection) were inserted into the left lateral ventricle (coordinates for intracerebroventricular injection (0° angle): anteroposterior, -0.5 mm; mediolateral, +1 mm; dorsoventral, -2 mm). The inserted guide cannula was fixed with dental cement to the skull, and a dummy cannula (Protech International, C315DCS-5, 3 mm guide, with 0.5 mm projection) was inserted into the guide cannula to avoid clogging. After a one-week recovery period, severe hypoglycemia was induced. For the glucose intracerebroventricular supplementation experiment, glucose dissolved in saline (100 μg/μl) or saline alone as control was intracerebroventricularly injected at a rate of 0.5 μl/min for 2 min, 0.5 and 3.5 hours after the first insulin injection. The time points for glucose intracerebroventricular injection were determined based on cerebrospinal fluid turnover rate, and the amount of glucose administered was selected to achieve cerebrospinal fluid glucose levels within the normal physiological range^46^. For the IL-1ra intracerebroventricular injection experiment, IL-1ra (1 μl/μg, administered at 0.5 μl/min for 6 min) or saline was intracerebroventricularly injected 1 hour prior to the intraperitoneal injection of 25% glucose. Four days after hypoglycemia induction, the same amount of IL-1ra or vehicle was intracerebroventricularly injected again.

### Stereotaxic RSC cannulation and drug delivery

The mice were anesthetized with a mixture of Zoletil (Virbac) and Rompun (Elanco), and 26-gauge guide cannulas (Protech International, C315GS-5/SPC, cut 2 mm below the pedestal) were implanted into the RSC (anteroposterior, -1.8 mm; mediolateral, -0.4 mm; dorsoventral, -0.75 mm (relative to bregma)). Dummy cannulas (Protech International, C315DCS-5/SPC, 0.5 mm projection) were kept in the guide cannulas until the experiment began. Mdivi-1 (10 μM), IL-1ra (1 mg/ml), or saline was initially injected into the RSC at a volume of 1 μl, 1 hour prior to the injection of 25% glucose and on the fourth day after hypoglycemia induction. All drugs were infused at a rate of 0.5 μl/min.

### *In vivo* siRNA silencing

For siRNA-mediated gene knockdown, siRNAs targeting *Dnm1l* and *Il1b*, as well as a negative control siRNA (Bioneer, SDH-1001), were reconstituted to a final concentration of 0.1 nmol/μl (10 nmol total) with RNase-free water. To facilitate delivery into the RSC, siRNA was complexed with Invivofectamine™ 3.0 reagent (Thermo Fisher Scientific, IVF3001) according to the manufacturer’s instructions. Briefly, siRNA at 100 pmol/μl was mixed 1:1 with the provided complexation buffer, followed by addition of Invivofectamine™ 3.0 reagent and incubation at 50°C for 30 minutes. Mice were anesthetized, and bilateral microinjections were performed targeting the RSC with stereotaxic surgery (anteroposterior, -1.8 mm; mediolateral, -0.4 mm; dorsoventral, -0.75 mm (relative to bregma)). The right hemisphere received *Dnm1l*- or *Il1b*-targeting siRNA, while the contralateral side received negative control siRNA. Each injection delivered 1 μl of siRNA solution (0.44 μg/μl) over 5 minutes, followed by an additional 5-minute dwell time to minimize reflux. The hypoglycemia experiment was conducted three days post-injection, and mice were sacrificed either 1 or 7 days after HPG.

### Fluorodeoxyglucose (FDG) PET/CT image acquisition

To acquire [^18^F]-FDG PET/CT images from each mouse (*n* = 5 per group) in the two groups (control and hypoglycemia), each mouse was fasted overnight after recovery from hypoglycemia before image acquisition. A heating pad was placed on the bed to maintain body temperature during the PET/CT scan. The head was fixed with a band to minimize movement. Static PET/CT images were obtained 60 minutes after a bolus injection of [^18^F]-FDG (12.21 ± 0.42 MBq in 0.2 mL saline) into a lateral tail vein. During image acquisition, mice were anesthetized with isoflurane (2.5% flow rate). Whole brain CT images were acquired using a micro-PET/CT scanner (nanoPET/CT, Bioscan). For CT image acquisition, the X-ray source was set to 200 μA and 45 kVp with a 0.5 mm filter. CT images were reconstructed using cone beam reconstruction with a Shepp filter with the cutoff at the Nyquist frequency and a binning factor of 4, resulting in an image matrix of 480 × 480 × 632 and a voxel size of 125 μm. For each PET scan, 44 transaxial images (43 × 32 pixel; 0.6 mm pixel size; 0.6 mm plane thickness) were reconstructed with an ordered subset using an expectation maximization iterative algorithm (4 iterations, 3 subsets).

### FDG PET/CT image processing

Cropping and template-based rigid co-registration into mouse brain PET template (created by the laboratory itself, 0.2 × 0.2 × 0.2 mm) space were performed using a PMOD medical image analysis software (PMOD Technologies LLC, RRID: SCR_016547, v4.1, Zurich, Swiss). An isotropic Gaussian filter with 0.4 mm full width at half maximum (FWHM) and reference count normalization were applied to generate a standardized uptake value ratio (SUVR) map for comparability between images. In this study, whole brain was used as the reference region.

### Nuclei extraction and snRNA-seq

For nuclei extraction, frozen samples were thawed at 4°C and kept on ice. Fresh and thawed samples were transferred into a precooled 1.5 ml tube and 1,000 µl of homogenization buffer (0.2 U µl^−1^ SUPERase In RNase Inhibitor (Thermo Fisher Scientific, cat. no. AM2694), 0.10% (v/v) Triton X-100, 1 µM DTT, 250 mM sucrose, 25 mM KCl, 5 mM MgCl2, 10 mM Tris-HCl, pH 8 in nuclease-free water) was added. The sample was pipetted several times and incubated on ice for 10 min. The sample was filtered through a 40 µm Flowmi Cell Strainer (Merck, cat. no. BAH136800040) to exclude larger debris. The nuclei suspension was centrifuged at 500*g* for 5 min at 4°C in a centrifuge to obtain a pellet of nuclei. After removing supernatant, the pellet was resuspended in 1% bovine serum albumin (BSA) in PBS with 0.2 U µl^−1^ SUPERase In RNase Inhibitor. Dissociated cells and extracted nuclei were suspended in 0.04% BSA in PBS and 1% BSA in PBS, respectively, containing 0.2 U µl^−1^ SUPERase RNase Inhibitor. Each sample was treated with either ReadyCount Green/Red Viability Stain (Invitrogen) for dissociated cells or YOYO-1 Iodide (Thermo Fisher Scientific) for nuclei. The number of cells per nuclei was measured, and cell viability was checked using the automated cell counter Countess 3 FL (Thermo Fisher Scientific). The cell/nuclei suspension was centrifuged at 400*g* for 5 min at 4°C and then resuspended in the BSA solution at an appropriate volume for microfluidic chip loading. Each sample was loaded per lane of a 10x microfluidic chip device Chromium Next GEM Chip G (10x Genomics). Approximately 16,000 nuclei were loaded per lane to target recovery of ∼10,000 nuclei per library.

### Single nucleus RNA sequencing analysis

Quality control was performed following ambient RNA correction and doublet identification described in the Main Methods. Low-quality nuclei were identified using median absolute deviation (MAD)–based filtering; nuclei exceeding 7 MADs from the median for library size or gene complexity were removed. Mitochondrial RNA–rich cells were additionally removed if their *pct_counts_mt* values exceeded 10% and fell outside 5 MADs of the median. Potential doublets were classified using Solo (v.1.3). After filtering, data integration was performed using scanpy (v.1.9.8) and scVI-tools (v.1.1.1). Highly variable genes (HVGs) were selected prior to integration using the seurat_v3 flavor with the top 5,000 HVGs identified from the raw count layer. For downstream integration, the dataset was prepared using scvi.setup_anndata and a variational autoencoder model was then trained with early-stopping based optimization. The trained SCVI latent space was subsequently used for clustering, visualization, and differential expression analyses. For batch-level comparisons, pseudobulk profiles were generated with decoupler (v.2.1.1). Differentially Expressed Genes (DEGs) were identified by using thresholds of |log2 fold change| ≥ 1 and P.value ≤ 0.05. Functional enrichment analysis of DEGs was performed using the web-based platforms Enrichr (https://maayanlab.cloud/Enrichr) and Metascape (https://metascape.org).

### Software and algorithms

|  |  |  |
| --- | --- | --- |
| Cellranger | https://www.10xgenomics.com/support/software/cell-ranger/latest | 10x Genomics Cell Ranger v9.0 |
| Cellbender | <sup>47</sup> | v.0.3.0 |
| Scanpy | <sup>48</sup> | v.1.9.8 |
| Solo | <sup>49</sup> | v.1.3 |
| scVI | <sup>50</sup> | v.1.1.1 |
| Decoupler | <sup>51</sup> | v.2.1.1 |
| Enrichr | <sup>52</sup> | https://maayanlab.cloud/Enrichr |
| Metascape | <sup>53</sup> | https://metascape.org |

### Immunohistochemistry

The mice were transcardially perfused with phosphate-buffered saline (PBS), and the brains were post-fixed overnight in 4% paraformaldehyde at 4°C. Subsequently, the brains were incubated in 30% sucrose at 4°C for 3 days, embedded in FSC 22 frozen section media (Leica Biosystems, 3801480), and sectioned at 25 μm using a cryostat (Thermo Fisher Scientific, Epredia™ Shandon™ Cryostat, CryoStar NX50). For antigen retrieval, brain slices were boiled at 95°C for 10 minutes in sodium citrate buffer (0.01 M, pH 6.0). After being washed in PBS, the brain sections were blocked with 0.3% Triton X-100 and 5% donkey serum in PBS for 1 hour at room temperature, followed by 24 hours of incubation at 4°C with primary antibodies [anti-4-HNE antiserum (Alpha Diagnostic, HNE11S, 1:200), anti-MAP2 (Millipore Sigma, AB5622, 1:200), anti-Iba-1 (Wako Chemicals, #01919741, 1:500; Abcam, ab5076, 1:500), anti-NeuN (EMD Millipore, MAB377, 1:200), anti-GFAP (Invitrogen, MA5-12023, 1:400), and anti-pDrp1 (S616) (Cell Signaling Technology, #3455, 1:100)]. The next day, after four washes in PBS, the slides were incubated with secondary antibodies [Alexa Fluor 488-conjugated donkey anti-rabbit (Invitrogen, A21206, 1:2,000), Alexa Fluor 568-conjugated donkey anti-rabbit (Invitrogen, A10042, 1:2,000), Alexa Fluor 594-conjugated donkey anti-mouse (Thermo Fisher Scientific, A21203, 1:2,000), and Alexa Fluor 647-conjugated donkey anti-goat (Thermo Fisher Scientific, A21447, 1:2,000)] for 2 hours at room temperature. After several washes, the slides were incubated with DAPI (0.5 μg/ml, Sigma‒Aldrich, D9542) for 5 minutes at room temperature and mounted with fluorescent mounting medium (DAKO, S3023). Images were obtained using a ZEISS LSM980 Airyscan2 confocal microscope and analyzed with ImageJ software (version 1.54d).

For unbiased screening of hypoglycemia-induced damage across the cortex, brain slice tissues from -1.3 mm to -3.64 mm from bregma were analyzed. We screened regions of the cortex in this area because topographic EEG mapping during severe hypoglycemia (30–50 mg/dL glucose) in patients with diabetes mellitus has revealed that overall changes in EEG are predominant in centrotemporal to parieto-occipital regions, suggesting that altered neuronal activity in these areas may be associated with hypoglycemia-induced damage^54^. Images were obtained using a ZEISS Axioscan 7 slide scanner.

### Western blot analysis

Dissected mouse RSC tissues were homogenized in lysis buffer containing RIPA lysis and extraction buffer (Thermo Scientific, 89900), a phosphatase inhibitor cocktail (Roche, PhosSTOP™, 4906845001) and a protease inhibitor cocktail (Roche, 04693159001). After the lysates were centrifuged for 20 min at 15,000*g*, the protein concentration in the supernatant was determined using the BCA Protein Assay Kit (Thermo Scientific, 23228). Equal amounts of protein were separated on 8% or 10% sodium dodecyl sulfate‒polyacrylamide gels and transferred to polyvinylidene difluoride membranes (Millipore, IPVH00010). The membranes were blocked with 5% skim milk for one hour at room temperature, followed by overnight incubation at 4°C with primary antibodies [anti-IL-1β (Proteintech, 16806-1-AP, 1:500), anti-TNF-α (Abcam, ab66579, 1:500), anti-IL-6 (Cell Signaling Technology, 12912S, 1:500), anti-Drp1 (Santa Cruz Biotechnology, SC-271583, 1:1,000), anti-phospho-Drp1 (Ser616) (Cell Signaling Technology, 3455S, 1:1,000), anti-PSD95 (Cell Signaling Technology, 3450S, 1:1,000), anti-synaptophysin (Sigma‒Aldrich, S5768, 1:500), anti-Mfn1 (ABclonal, A9880, 1:1,000), anti-Mfn2 (ABclonal, A19678, 1:1,000), anti-OPA1 (ABclonal, A9833, 1:1,000), anti-VDAC1 (Abcam, ab14734, 1:1,000), and anti-β-actin (Santa Cruz Biotechnology, SC-81178, 1:3,000)]. After three washes, the membranes were incubated with horseradish peroxidase (HRP)-conjugated secondary antibodies [goat anti-mouse (Invitrogen, 31430, 1:2,000) and goat anti-rabbit (Invitrogen, 31460, 1:2,000)] for one hour at room temperature. The membranes were visualized with enhanced chemiluminescence (ECL) substrate (Thermo Scientific, 32106) using an Amersham Imager 600 instrument (GE Healthcare, Chicago, Illinois, USA), and the signals were quantified using ImageJ software.

### Mitochondria isolation

Mitochondria were isolated from fresh retrosplenial cortex (RSC) tissue at day 1 after hypoglycemia experiment using the Mitochondria Isolation Kit for Tissue (Thermo Scientific, 89801) according to the manufacturer’s instructions. Briefly, dissected RSC tissues were immediately placed in ice-cold isolation buffer to maintain mitochondrial integrity. The tissue was homogenized in Mitochondria Isolation Reagent A using a Dounce homogenizer. The resulting homogenate was subjected to differential centrifugation at 4°C to separate the mitochondrial fraction from the cytosolic and nuclear components. Following isolation, the mitochondrial pellet was solubilized in RIPA lysis and extraction buffer (Thermo Scientific, 89900).

### TUNEL assay

The TUNEL assay was performed using the In Situ Cell Death Detection Kit, Fluorescein (Roche, #11684795910), which was utilized to detect apoptosis-related DNA fragmentation. Staining was performed according to the manufacturer’s instructions. For antigen retrieval, brain slices were boiled at 95°C for 10 minutes in sodium citrate buffer (0.01 M, pH 6.0). After three washes with PBS, the slides were incubated with a TUNEL reaction mixture, which contained terminal deoxynucleotidyl transferase (TdT) and fluorescein-dUTP, under humid conditions for 1 hour at 37°C. Following several washes, the slides were incubated with DAPI for 5 minutes at room temperature and mounted with fluorescent mounting medium (DAKO, S3023). Images were captured using a ZEISS LSM980 Airyscan2 confocal microscope and analyzed using ImageJ software.

### Transmission electron microscopy (TEM)

The mice were sacrificed, and their brains were fixed for 18 hours with 2.5% glutaraldehyde (Agar Scientific, AGR1554) in 0.1 M phosphate buffer (pH 7.4). TEM images of the RSC were taken at 15,000x magnification with a JEM-1400 (JEOL, Japan). Images of randomly selected areas of the RSC were captured by an operator who was blinded to the groups. Mitochondrial analysis was conducted using ImageJ software.

### Microglial morphology analysis

Z-stack confocal images of Iba-1-stained microglia in the RSC were acquired at 63x magnification, with a 25 μm thickness and an interval of 1 μm. Images were captured using a ZEISS LSM980 Airyscan2 confocal microscope. The images were then reconstructed with the Simple Neurite Tracer plug-in of Fiji software^55^. Microglia were manually traced by selecting each branch point to the border, and the software traced the path between these points.

### Cell culture and Transwell co-culture system

BV-2 murine microglia and SH-SY5Y human neuronal cells were cultured in Dulbecco’s modified Eagle’s medium (DMEM, Welgene, Gyeongsan, Korea, LM001-05) supplemented with 10% fetal bovine serum (FBS, Gibco, # 16000044) and 1% penicillin/streptomycin (Welgene, LS 202-02). The cells were maintained in a humidified incubator containing 5% CO2 at 37°C. Before glucose deprivation, the SH-SY5Y cells were differentiated using 50 µM retinoic acid (Sigma‒ Aldrich, R2625) for 5 days.

For co-culture experiments, SH-SY5Y or BV-2 cells were plated on either 6-well plates or the upper chamber of Transwell inserts (Corning, CLS3450). For mdivi-1, minocycline or IL-1ra pretreatment, before glucose deprivation, SH-SY5Y or BV-2 cells were individually treated with mdivi-1 (25 µM), minocycline (50 µM), or IL-1ra (1 µg/ml) for 6 hours. After drug pretreatment, all the plates were washed with Dulbecco’s phosphate-buffered saline (DPBS), and the medium was changed to glucose-free DMEM (Welgene, LM001-79) supplemented with 10% FBS and 1% penicillin/streptomycin for 24 hours. The control groups received the same volume of vehicle (DMSO or ethanol). When the medium was changed, Transwell inserts were placed into the 6-well plates for co-culture experiments with glucose-deprived medium.

### Lactate dehydrogenase (LDH) assay

To measure cell cytotoxicity, the amount of released LDH was analyzed using an LDH cytotoxicity assay kit (DoGenBio, DG-LDH500). After glucose deprivation for 24 hours, the medium from each well was transferred to a 96-well plate and incubated with LDH reaction mixture at 36°C in the dark. The absorbance was measured at 450 nm using a microplate reader (Tecan, Infinite m200 Pro). The percentage of cytotoxicity was calculated according to the manufacturer’s instructions.

### Immunocytochemistry

SH-SY5Y cells or BV-2 cells were seeded on poly-D-lysine (Sigma‒Aldrich, P7280)-coated glass coverslips. After 24 hours of glucose deprivation, the cells were washed twice with PBS and then fixed with 4% paraformaldehyde for 30 minutes at room temperature. After being washed, the cells were permeabilized and blocked with 1% bovine serum albumin (BSA) and 0.2% Triton X-100 in PBS for 1 hour at room temperature, followed by overnight incubation at 4°C with anti-cleaved caspase-3 (Cell Signaling Technology, 9664S, 1:150) and anti-CD68 (Bio-Rad, MCA1957GA, 1:200) antibodies. The next day, after three washes, the cells were incubated with secondary antibody for 2 hours at room temperature. After being washed, the cells were stained with DAPI for 5 minutes at room temperature and mounted with fluorescent mounting medium (DAKO, S3023). The cells were visualized using a ZEISS LSM980 Airyscan2 confocal microscope and analyzed with ImageJ software.

### MitoTracker staining and mitochondrial length analysis

After the *in vitro* experiments, SH-SY5Y cells were incubated with MitoTracker Red CMXRos (Thermo Scientific, M7512, 100 nM) for 30 minutes and then fixed. Z-stack confocal images at intervals of 0.25 μm were acquired at 63x magnification. Images were captured using a ZEISS LSM980 Airyscan2 confocal microscope. The length of the major mitochondrial axis was quantified with the Simple Neurite Tracer plug-in of Fiji software. In brief, each end point of the major mitochondrial axis was manually selected, and then the software traced the entire mitochondrial axis to measure its length.

### Morris water maze test

The mice were trained in a 120 cm diameter circular pool filled with water at a height of 35 cm. The water was rendered opaque using nontoxic white paint and maintained at 25°C. The movement of the mice was recorded with a video camera suspended over the central area of the pool. The escape platform was submerged 1 cm below the water level in the middle of the southeast quadrant. During the acquisition phase, each mouse underwent four training trials daily for five consecutive days. The mice were released from different starting points and allowed to search for the hidden platform for a maximum of 90 seconds. If a mouse found the platform within 90 seconds, it was allowed to stay on the platform for 30 seconds. If a mouse could not find the platform within 90 seconds, it was manually guided to the platform. After the training session for the learning phase, a probe trial and retrieval trial were conducted to assess spatial memory consolidation^56^. The probe test, in which the platform was removed from the target quadrant, was conducted three days after the last acquisition trial. Each mouse was allowed to swim for 60 seconds. Seven days after the probe trial, the retrieval test was conducted to assess memory retrieval. The time spent in the target quadrant and the mean distance required to reach the platform were measured with EthoVision XT 14 software.

### Open field test

White acrylic boxes were employed for the open field test (40 × 40 cm). Each mouse was individually placed in the center zone of the apparatus and allowed to explore freely for 10 minutes. The movements of the mice were recorded. The center zone was defined as a 20 × 20 cm area in the middle of the apparatus. The data were analyzed using EthoVision XT 14 software.

### Elevated plus maze

The elevated plus maze consisted of two opposite open arms and two closed arms (each 30 × 6 cm) and was elevated 60 cm above the floor. Each mouse was placed in the center of the maze and allowed to move freely for 5 minutes. Their movements were recorded, and the time spent in the open arms was manually measured.

### Tail suspension test

Each mouse was suspended by the tip of its tail with adhesive tape from an acrylic column positioned 35 cm above the floor. The behavior of each mouse was observed for 5 minutes. Movements were recorded by a camera, and the immobility time was manually measured. The immobility time was defined as the amount of time during which the mouse did not move for more than 1 second.

### Statistics

All the animals were randomly grouped within each batch, and the cells were randomly assigned to the experimental groups. No data were excluded from the analyses, and the investigators were not blinded to the drug treatment conditions. IBM SPSS v26.0 software (SPSS Inc., Chicago, Illinois, USA) was used to perform statistical analysis. Statistical significance was determined by an unpaired two-tailed Student’s *t* test, multiple *t* tests with the Holm‒Sidak method, or analysis of variance (ANOVA) (one-way or two-way) followed by Tukey’s or Dunnett’s post hoc test. The data are shown as the means ± SEMs, and *P* < 0.05 was considered to indicate statistical significance.

For FDG PET/CT image analysis, each parametric image was spatially normalized to a mouse brain MRI template. Image analyses were performed using SnPM (Wellcome Trust Centre for Neuroimaging). The registered images were subjected to statistical analysis using a t-test for group comparisons. Comparisons between the test and vehicle groups were conducted using nonparametric permutation tests. A corrected p-value threshold of < 0.05 was applied for statistical significance.

## Acknowledgments

We thank Prof. Sang Won Suh and Dr. A Ra Kho (Hallym Univ.) for sharing valuable tips to setup mouse model of hypoglycemia. We thank Prof. Min-Seon Kim (Ulsan Univ.) for critical questions and discussion. We thank Prof. Myoung-Hwan Kim and Woo Seok Song (Seoul National Univ.) for technical assistance, and Prof. Seung-Jae Lee and Prof. Eun-Hee Shin (Seoul National Univ.) for kindly providing SH-SY5Y cells and BV-2 cells, respectively. We thank Dr. Ji Young Mun (Korea Brain Research Institute) for the advice on processing TEM data. The schematics in the figures were created using BioRender.

## Author contributions

O.K. conceived the study concept and design; J.-Y.J., S.L., M.K.S., and S.K. performed experiments and processed samples; K.P. and J.-I.K constructed scRNA libraries; M.K., H. L., J.-I.K. performed data analysis of snRNA-seq; D.K. and H.-Y.L. performed FDG-PET; J.-Y.J., S.L., M.K.S., S.P., J.H.H., and O.K. analyzed and contributed to visualization of the data; J.-Y.J., S.L., and O.K. interpreted the results and wrote the manuscript. J.-Y.J. and S.L. contributed equally to the work. All authors edited, reviewed, and approved the final version of the manuscript for publication. O.K. is the guarantors for this paper.

## Data and materials availability

Transcriptomic datasets from this manuscript have been uploaded to the NCBI Gene Expression Omnibus (GEO: GSE314246). This paper does not report any original codes. Any additional information required to reanalyze the data reported in this paper is available from the lead contact upon request.

## References and Notes

1 American Diabetes Association Professional Practice Committee for Diabetes. 9. Pharmacologic Approaches to Glycemic Treatment: Standards of Care in Diabetes-2026. Diabetes Care 49, S183–s215 (2026). 10.2337/dc26-S009

2 Cho, N. H., et al. Patient Understanding of Hypoglycemia in Tertiary Referral Centers. Diabetes Metab J 42, 43–52 (2018). 10.4093/dmj.2018.42.1.43

3 Suh, S. W., Aoyama, K., Matsumori, Y., Liu, J. & Swanson, R. A. Pyruvate administered after severe hypoglycemia reduces neuronal death and cognitive impairment. Diabetes 54, 1452–1458 (2005). 10.2337/diabetes.54.5.1452

4 Graveling, A. J., Deary, I. J. & Frier, B. M. Acute hypoglycemia impairs executive cognitive function in adults with and without type 1 diabetes. Diabetes Care 36, 3240–3246 (2013). 10.2337/dc13-0194

5 Nishihama, K., et al. Sudden Death Associated with Severe Hypoglycemia in a Diabetic Patient During Sensor-Augmented Pump Therapy with the Predictive Low Glucose Management System. Am J Case Rep 22, e928090 (2021). 10.12659/ajcr.928090

6 Basu, S., et al. Estimation of global insulin use for type 2 diabetes, 2018-30: a microsimulation analysis. Lancet Diabetes Endocrinol 7, 25–33 (2019). 10.1016/s2213-8587(18)30303-6

7 Zhong, V. W., et al. Incidence and Trends in Hypoglycemia Hospitalization in Adults With Type 1 and Type 2 Diabetes in England, 1998-2013: A Retrospective Cohort Study. Diabetes Care 40, 1651–1660 (2017). 10.2337/dc16-2680

8 Youle, R. J. & van der Bliek, A. M. Mitochondrial fission, fusion, and stress. Science 337, 1062–1065 (2012). 10.1126/science.1219855

9 Ni, H. M., Williams, J. A. & Ding, W. X. Mitochondrial dynamics and mitochondrial quality control. Redox Biol 4, 6–13 (2015). 10.1016/j.redox.2014.11.006

10 Liesa, M. & Shirihai, O. S. Mitochondrial dynamics in the regulation of nutrient utilization and energy expenditure. Cell Metab 17, 491–506 (2013). 10.1016/j.cmet.2013.03.002

11 Becher, B., Spath, S. & Goverman, J. Cytokine networks in neuroinflammation. Nat Rev Immunol 17, 49–59 (2017). 10.1038/nri.2016.123

12 Heneka, M. T., et al. Neuroinflammation in Alzheimer’s disease. Lancet Neurol 14, 388–405 (2015). 10.1016/s1474-4422(15)70016-5

13 Xu, Y., et al. The effect of lithium chloride on the attenuation of cognitive impairment in experimental hypoglycemic rats. Brain Res Bull 149, 168–174 (2019). 10.1016/j.brainresbull.2019.04.019

14 Auer, R. N., Wieloch, T., Olsson, Y. & Siesjö, B. K. The distribution of hypoglycemic brain damage. Acta Neuropathol 64, 177–191 (1984). 10.1007/bf00688108

15 Ennis, K., Tran, P. V., Seaquist, E. R. & Rao, R. Postnatal age influences hypoglycemia-induced neuronal injury in the rat brain. Brain Res 1224, 119–126 (2008). 10.1016/j.brainres.2008.06.003

16 Taguchi, N., Ishihara, N., Jofuku, A., Oka, T. & Mihara, K. Mitotic phosphorylation of dynamin-related GTPase Drp1 participates in mitochondrial fission. J Biol Chem 282, 11521–11529 (2007). 10.1074/jbc.M607279200

17 Hanisch, U. K. Microglia as a source and target of cytokines. Glia 40, 140–155 (2002). 10.1002/glia.10161

18 Verdonk, F., et al. Phenotypic clustering: a novel method for microglial morphology analysis. J Neuroinflammation 13, 153 (2016). 10.1186/s12974-016-0614-7

19 Todd, T. P. & Bucci, D. J. Retrosplenial Cortex and Long-Term Memory: Molecules to Behavior. Neural Plast 2015, 414173 (2015). 10.1155/2015/414173

20 Sestieri, C., Corbetta, M., Romani, G. L. & Shulman, G. L. Episodic memory retrieval, parietal cortex, and the default mode network: functional and topographic analyses. J Neurosci 31, 4407–4420 (2011). 10.1523/jneurosci.3335-10.2011

21 Hershey, T., et al. Frequency and timing of severe hypoglycemia affects spatial memory in children with type 1 diabetes. Diabetes Care 28, 2372–2377 (2005). 10.2337/diacare.28.10.2372

22 Wright, R. J., Frier, B. M. & Deary, I. J. Effects of acute insulin-induced hypoglycemia on spatial abilities in adults with type 1 diabetes. Diabetes Care 32, 1503–1506 (2009). 10.2337/dc09-0212

23 Broadbent, N. J., Squire, L. R. & Clark, R. E. Spatial memory, recognition memory, and the hippocampus. Proc Natl Acad Sci U S A 101, 14515–14520 (2004). 10.1073/pnas.0406344101

24 Alexander, A. S., Place, R., Starrett, M. J., Chrastil, E. R. & Nitz, D. A. Rethinking retrosplenial cortex: Perspectives and predictions. Neuron 111, 150–175 (2023). 10.1016/j.neuron.2022.11.006

25 Ash, J. A., et al. Functional connectivity with the retrosplenial cortex predicts cognitive aging in rats. Proc Natl Acad Sci U S A 113, 12286–12291 (2016). 10.1073/pnas.1525309113

26 Wu, Y. T., et al. Quantitative relationship between cerebrovascular network and neuronal cell types in mice. Cell Rep 39, 110978 (2022). 10.1016/j.celrep.2022.110978

27 Bolo, N. R., et al. Functional Connectivity of Insula, Basal Ganglia, and Prefrontal Executive Control Networks during Hypoglycemia in Type 1 Diabetes. J Neurosci 35, 11012–11023 (2015). 10.1523/jneurosci.0319-15.2015

28 Bolo, N. R., et al. Brain activation during working memory is altered in patients with type 1 diabetes during hypoglycemia. Diabetes 60, 3256–3264 (2011). 10.2337/db11-0506

29 Nong, H., et al. Alterations in intra- and inter-network connectivity associated with cognition impairment in insulinoma patients. Front Endocrinol (Lausanne) 14, 1234921 (2023). 10.3389/fendo.2023.1234921

30 Dong, W. T., et al. Mitochondrial fission drives neuronal metabolic burden to promote stress susceptibility in male mice. Nat Metab 5, 2220–2236 (2023). 10.1038/s42255-023-00924-6

31 Hajimohammadreza, I., et al. A specific inhibitor of calcium/calmodulin-dependent protein kinase-II provides neuroprotection against NMDA- and hypoxia/hypoglycemia-induced cell death. J Neurosci 15, 4093–4101 (1995). 10.1523/jneurosci.15-05-04093.1995

32 Li, Y. H. & Gong, P. L. Neuroprotective effect of dauricine in cortical neuron culture exposed to hypoxia and hypoglycemia: involvement of correcting perturbed calcium homeostasis. Can J Physiol Pharmacol 85, 621–627 (2007). 10.1139/y07-056

33 Zhao, L., Li, S., Wang, S., Yu, N. & Liu, J. The effect of mitochondrial calcium uniporter on mitochondrial fission in hippocampus cells ischemia/reperfusion injury. Biochem Biophys Res Commun 461, 537–542 (2015). 10.1016/j.bbrc.2015.04.066

34 Won, S. J., et al. Prevention of hypoglycemia-induced neuronal death by minocycline. J Neuroinflammation 9, 225 (2012). 10.1186/1742-2094-9-225

35 Bergold, P. J., Furhang, R. & Lawless, S. Treating Traumatic Brain Injury with Minocycline. Neurotherapeutics 20, 1546–1564 (2023). 10.1007/s13311-023-01426-9

36 Zhao, F., Hua, Y., He, Y., Keep, R. F. & Xi, G. Minocycline-induced attenuation of iron overload and brain injury after experimental intracerebral hemorrhage. Stroke 42, 3587–3593 (2011). 10.1161/strokeaha.111.623926

37 Plane, J. M., Shen, Y., Pleasure, D. E. & Deng, W. Prospects for minocycline neuroprotection. Arch Neurol 67, 1442–1448 (2010). 10.1001/archneurol.2010.191

38 Scott, G., et al. Minocycline reduces chronic microglial activation after brain trauma but increases neurodegeneration. Brain 141, 459–471 (2018). 10.1093/brain/awx339

39 Boutin, H., et al. Role of IL-1alpha and IL-1beta in ischemic brain damage. J Neurosci 21, 5528–5534 (2001). 10.1523/jneurosci.21-15-05528.2001

40 Craft, J. M., Watterson, D. M., Hirsch, E. & Van Eldik, L. J. Interleukin 1 receptor antagonist knockout mice show enhanced microglial activation and neuronal damage induced by intracerebroventricular infusion of human beta-amyloid. J Neuroinflammation 2, 15 (2005). 10.1186/1742-2094-2-15

41 Lu, K. T., Wang, Y. W., Wo, Y. Y. & Yang, Y. L. Extracellular signal-regulated kinase-mediated IL-1-induced cortical neuron damage during traumatic brain injury. Neurosci Lett 386, 40–45 (2005). 10.1016/j.neulet.2005.05.057

42 Rothwell, N. Interleukin-1 and neuronal injury: mechanisms, modification, and therapeutic potential. Brain Behav Immun 17, 152–157 (2003). 10.1016/s0889-1591(02)00098-3

43 Harrison, J. K., et al. Role for neuronally derived fractalkine in mediating interactions between neurons and CX3CR1-expressing microglia. Proc Natl Acad Sci U S A 95, 10896–10901 (1998). 10.1073/pnas.95.18.10896

44 Butler, C. A., et al. Microglial phagocytosis of neurons in neurodegeneration, and its regulation. J Neurochem 158, 621–639 (2021). 10.1111/jnc.15327

45 Terstege, D. J., et al. Impaired parvalbumin interneurons in the retrosplenial cortex as the cause of sex-dependent vulnerability in Alzheimer’s disease. Sci Adv 11, eadt8976 (2025). 10.1126/sciadv.adt8976

46 Nigrovic, L. E., Kimia, A. A., Shah, S. S. & Neuman, M. I. Relationship between cerebrospinal fluid glucose and serum glucose. N Engl J Med 366, 576–578 (2012). 10.1056/NEJMc1111080

47 Fleming, S. J., et al. Unsupervised removal of systematic background noise from droplet-based single-cell experiments using CellBender. Nat Methods 20, 1323–1335 (2023). 10.1038/s41592-023-01943-7

48 Wolf, F. A., Angerer, P. & Theis, F. J. SCANPY: large-scale single-cell gene expression data analysis. Genome Biol 19, 15 (2018). 10.1186/s13059-017-1382-0

49 Bernstein, N. J., et al. Solo: Doublet Identification in Single-Cell RNA-Seq via Semi-Supervised Deep Learning. Cell Syst 11, 95–101.e105 (2020). 10.1016/j.cels.2020.05.010

50 Lopez, R., Regier, J., Cole, M. B., Jordan, M. I. & Yosef, N. Deep generative modeling for single-cell transcriptomics. Nat Methods 15, 1053–1058 (2018). 10.1038/s41592-018-0229-2

51 Badia, I. M. P., et al. decoupleR: ensemble of computational methods to infer biological activities from omics data. Bioinform Adv 2, vbac016 (2022). 10.1093/bioadv/vbac016

52 Kuleshov, M. V., et al. Enrichr: a comprehensive gene set enrichment analysis web server 2016 update. Nucleic Acids Res 44, W90–97 (2016). 10.1093/nar/gkw377

53 Zhou, Y., et al. Metascape provides a biologist-oriented resource for the analysis of systems-level datasets. Nat Commun 10, 1523 (2019). 10.1038/s41467-019-09234-6

54 Tribl, G., et al. EEG topography during insulin-induced hypoglycemia in patients with insulin-dependent diabetes mellitus. Eur Neurol 36, 303–309 (1996). 10.1159/000117277

55 Schindelin, J., et al. Fiji: an open-source platform for biological-image analysis. Nat Methods 9, 676–682 (2012). 10.1038/nmeth.2019

56 Vorhees, C. V. & Williams, M. T. Morris water maze: procedures for assessing spatial and related forms of learning and memory. Nat Protoc 1, 848–858 (2006). 10.1038/nprot.2006.116

