## Supplementary Information for "Retrosplenial cortex vulnerability links severe hypoglycemia to cognitive impairment through neuron–microglia crosstalk"

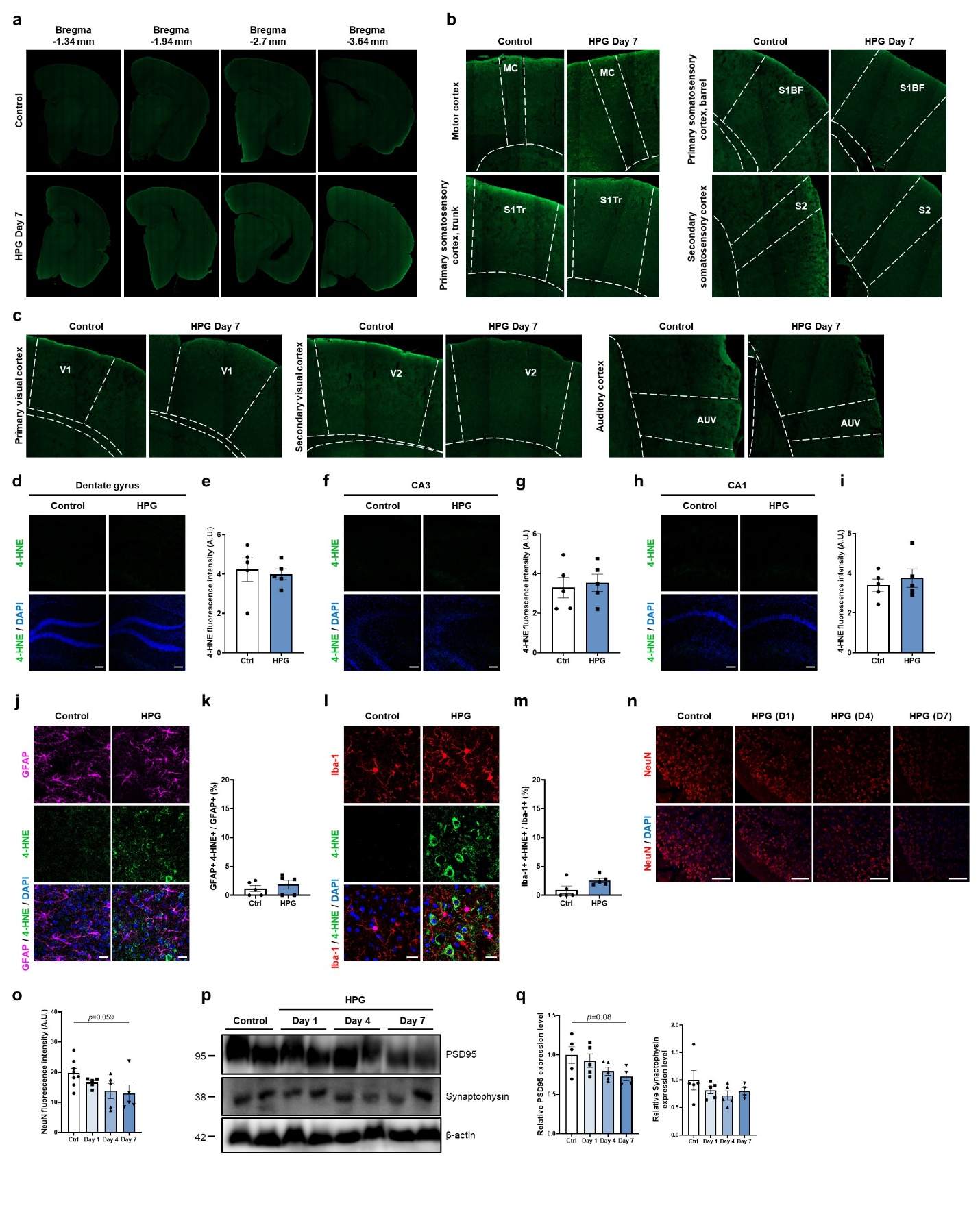

**Supplementary Fig. 1**. **Characterization of acute hypoglycemia-induced damage in the cortex and hippocampus. Related to Figure 1.** (a) Unbiased screening was conducted in the cortex region from Bregma -1.3 mm to Bregma -3.64 mm. (b and c) Representative confocal images with 4-HNE staining in the RSC, motor cortex (MC), primary somatosensory cortex trunk (S1Tr), primary somatosensory cortex barrel field (S1BF), secondary somatosensory cortex (S2), primary visual cortex (V1), secondary visual cortex (V2), and auditory cortex (AUV). (d) Representative confocal images of the dentate gyrus with 4-HNE staining from the control (Ctrl) and hypoglycemia-experienced (HPG) mice. Scale bars, 100 μm. (e) Quantification of 4-HNE intensity (*n* = 5 mice per group). (f) Representative confocal images of the CA3 with 4-HNE. Scale bars, 100 μm. (g) Quantification of 4-HNE intensity (*n* = 5 mice per group). (h) Representative confocal images of the CA1 with 4-HNE. Scale bars, 100 μm. (i) Quantification of 4-HNE intensity (*n* = 5 mice per group). (j) Representative confocal images of the RSC tissues with GFAP and 4-HNE co-staining in the control and hypoglycemic group (HPG) at day 7. Scale bars, 20 μm. (k) Quantification of the percentage of 4-HNE^+^ cells co-localized with GFAP^+^ cells (*n* = 5 mice per group). (l) Representative confocal images of the RSC tissues with Iba-1 and 4-HNE co-staining. Scale bars, 20 μm. (m) Quantification of the percentage of 4-HNE^+^ cells co-localized with Iba-1^+^ cells (*n* = 5 mice per group). (n) Representative confocal images of NeuN staining. Scale bars, 100 μm. (o) Quantification of NeuN intensity in the RSC region (*n* = 5 or 8 mice per group). (p) Representative immunoblots from the RSC tissues during the progression of hypoglycemic damage. PSD95 is a post-synaptic protein, and synaptophysin is a pre-synaptic vesicle protein. (q) Quantification of PSD95 (left) and synaptophysin (right) relative expression levels (*n* = 4 or 5 mice per group). Statistical analysis were performed using two-tailed Student’s *t* test (e, g, i, k and m) and one-way ANOVA followed by Dunnett’s multiple comparisons test (o and q). Data are presented as mean ± SEM.

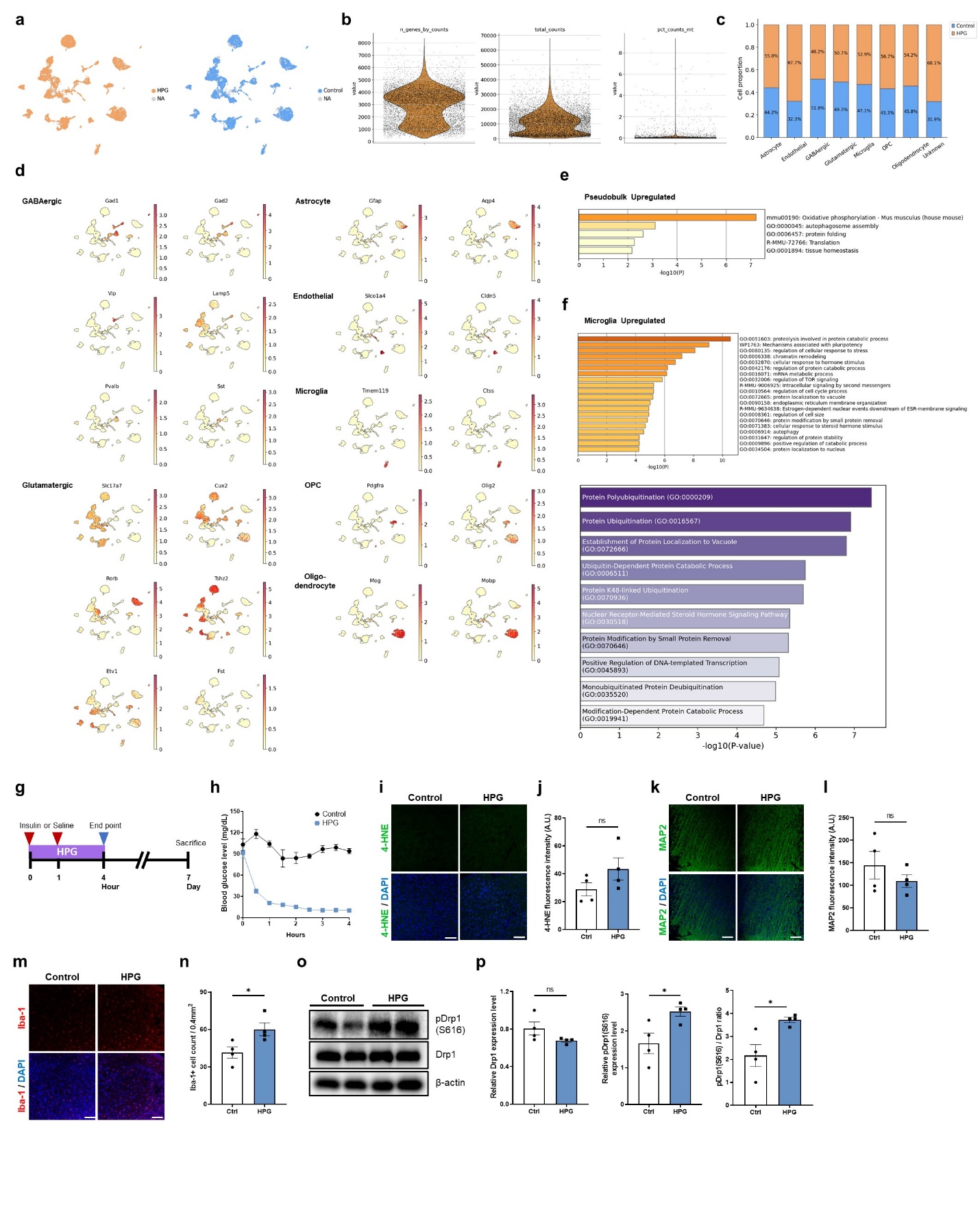

**Supplementary Fig. 2**. **snRNA-seq analysis and shortened hypoglycemia exposure model. Related to Figure 2.** (a) UMAP projection of all cells, split by sample. (b) Violin plots showing the distribution of detected features (n_genes_by_counts), total UMI counts (total_counts), and percentage of mitochondrial transcripts (pct_counts_mt) in the quality-controlled and integrated dataset. (c) Proportion of Control and HPG cells within each annotated cell type. (d) Expression of key cell type marker genes visualized on the UMAP. (e) Functional enrichment analysis of genes upregulated in HPG performed using Metascape. (f) Functional enrichment analysis of genes upregulated in HPG microglia compared with Control microglia using Metascape (up) and Enrichr (down). (g) Experimental design. C57BL/6 mice were administered insulin (HPG, *n* = 4 mice) or saline (Control, *n* = 4 mice) twice, with a one-hour interval. To terminate the hypoglycemia, 25% glucose was administered intraperitoneally four hours after the initial insulin administration. (h) Blood glucose levels (mg/dL) of the HPG and control group. Representative confocal images of the RSC tissues stained with (i) 4-HNE, (k) MAP2, and (m) Iba-1. Scale bar, 100 μm. Quantitative analysis of mean fluorescence intensity for (j) 4-HNE and (l) MAP2 in the control (Ctrl) and hypoglycemia (HPG) groups. (n) Quantitative analysis of the number of Iba-1 cells. (o) Representative immunoblots showing pDrp1(S616) and Drp1 expression. (p) Quantitative analysis of the relative expression levels of Drp1, pDrp1(S616), and the ratio of pDrp1(S616)/Drp1. Statistical analysis was performed using two-tailed Student’s *t* test (j, l, n, and p). Data are presented as mean ± SEM (HPG, *n* = 4 mice; Control, *n* = 4 mice). *P < 0.05; ns, not significant.

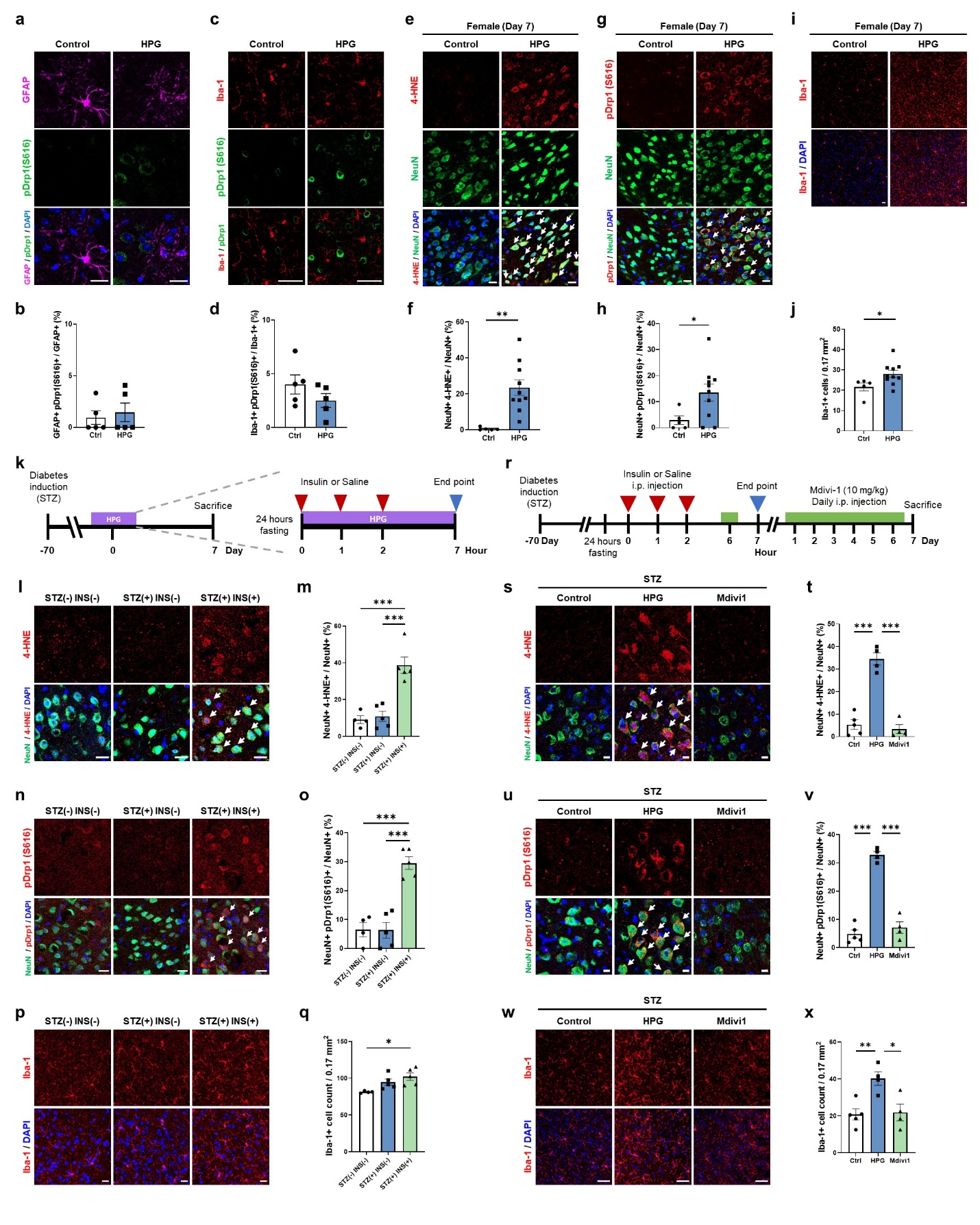

**Supplementary Fig. 3**. **Additional immunofluorescence data in male, female mice, and STZ-induced diabetic male mice. Related to Figure 3.** (a) Representative confocal images of the RSC tissues with GFAP and pDrp1(S616) co-staining. Scale bars, 20 μm. (b) Quantification of the percentage of pDrp1(S616)^+^ cells co-localized with GFAP^+^ cells (*n* = 5 mice per group). (c) Representative confocal images of the RSC tissues with Iba-1 and pDrp1(S616) co-staining. Scale bars, 50 μm. (d) Quantification of the percentage of pDrp1(S616)^+^ cells co-localized with Iba-1^+^ cells (*n* = 5 mice per group). (e) Representative confocal images of the RSC tissues with NeuN and 4-HNE co-staining in female mice. Scale bars, 20 μm. White arrows indicate 4-HNE^+^ cells co-localized with NeuN^+^ cells. (f) Quantification of the percentage of 4-HNE^+^ cells co-localized with NeuN^+^ cells (*n* = 5 or 10 mice per group). (g) Representative confocal images of the RSC tissues with NeuN and pDrp1 co-staining in female mice. Scale bars, 20 μm. White arrows indicate pDrp1^+^ cells co-localized with NeuN^+^ cells. (h) Quantification of the percentage of pDrp1(S616)^+^ cells co-localized with NeuN^+^ cells (*n* = 5 or 10 mice per group). (i) Representative confocal images of the RSC tissues with Iba-1 staining in female mice. Scale bars, 20 μm. (j) Quantification of Iba-1^+^ cells normalized by the area size of 0.17 mm^2^ (*n* = 5 or 10 mice per group). (k) Experimental design of the study. C57BL/6 mice were i.p. injected with vehicle or STZ (55 mg/kg) for five consecutive days. After diabetes induction (blood glucose level exceeding 200 mg/dL), the diabetic condition was maintained for 10 weeks prior to the hypoglycemia experiment. The following groups were analyzed: non-diabetic control group without insulin-induced hypoglycemia (STZ(-)INS(-), *n* = 4), diabetic control group without insulin-induced hypoglycemia (STZ(+)INS(-), *n* = 5), and diabetic group with insulin-induced hypoglycemia (STZ(+)INS(+), *n* = 5). (l) Representative confocal images of the RSC tissues with NeuN and 4-HNE co-staining. Scale bars, 20 μm. White arrows indicate 4-HNE^+^ cells co-localized with NeuN^+^ cells. (m) Quantification of the percentage of 4-HNE^+^ cells co-localized with NeuN^+^ cells. (n) Representative confocal images of the RSC tissues with NeuN and pDrp1(S616) co-staining. Scale bars, 20 μm. White arrows indicate pDrp1^+^ cells co-localized with NeuN^+^ cells. (o) Quantification of the percentage of pDrp1^+^ cells co-localized with NeuN^+^ cells. (p) Representative confocal images of the RSC tissues with Iba-1 staining. Scale bars, 20 μm. (q) Quantification of Iba-1^+^ cells normalized by the area size of 0.17 mm^2^. (r) Experimental design of the study. The diabetic control group without insulin-induced hypoglycemia (Control; *n* = 5) received vehicle injection, whereas diabetic mice subjected to insulin-induced hypoglycemia were treated with either vehicle (HPG; *n* = 4) or mdivi-1 (Mdivi-1; *n* = 4). (s) Representative confocal images of the RSC tissues with NeuN and 4-HNE co-staining. Scale bars, 10 μm. White arrows indicate 4-HNE^+^ cells co-localized with NeuN^+^ cells. (t) Quantification of the percentage of 4-HNE^+^ cells co-localized with NeuN^+^ cells. (u) Representative confocal images of the RSC tissues with NeuN and pDrp1(S616) co-staining. Scale bars, 10 μm. White arrows indicate pDrp1^+^ cells co-localized with NeuN^+^ cells. (v) Quantification of the percentage of pDrp1^+^ cells co-localized with NeuN^+^ cells. (w) Representative confocal images of the RSC tissues with Iba-1 staining. Scale bars, 50 μm. (x) Quantification of Iba-1^+^ cells normalized by the area size of 0.17 mm^2^. Statistical analysis were performed using two-tailed Student’s *t* test (b, d, f, h, and j) and one-way ANOVA followed by Tukey’s multiple comparison test (m, o, q, t, v, and x). Data are presented as means ± SEM. *P < 0.05; **P < 0.01; ***P < 0.001.

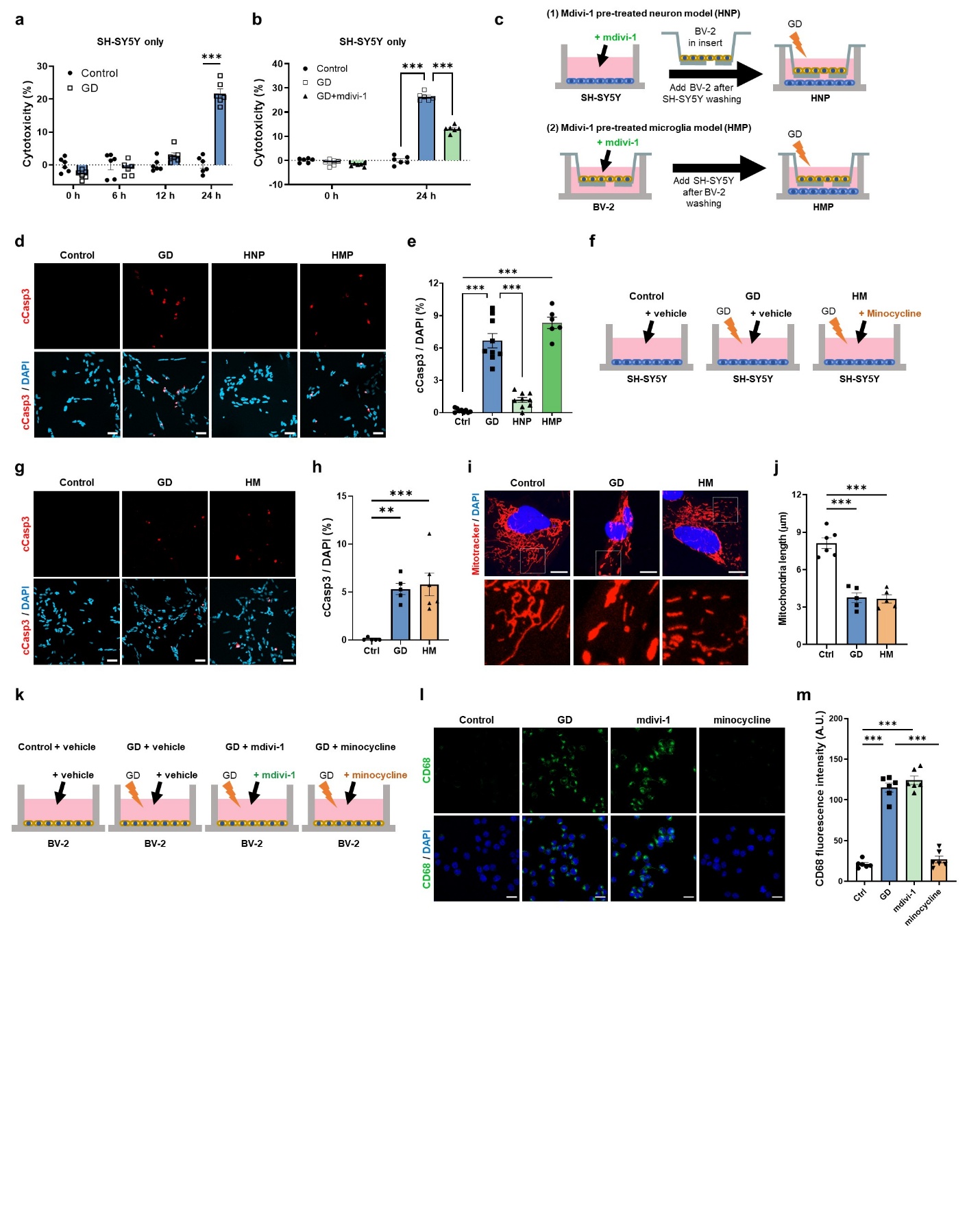

**Supplementary Fig. 4. Neither mdivi-1 directly prevent microglial activation, nor does minocycline prevent neuronal mitochondrial fission and apoptosis *in vitro*. Related to Figure 4.** (a) SH-SY5Y cells were previously incubated in normal glucose-containing media (25 mM), then changed to glucose-containing media (Control, 25 mM) or glucose-deprived media (GD, 0 mM) for the experiment. Cytotoxicity was quantified by LDH assay at 0, 6, 12, and 24 hours after the media change (*n* = 6 independent samples per group). (b) Glucose-containing media (25 mM) with vehicle (Control) or glucose-deprived media with vehicle (GD) or mdivi-1 (GD+mdivi-1, 25 μM) were treated in SH-SY5Y cells for 24 hours. Cytotoxicity was quantified by LDH assay at 24 hours after the media changed (*n* = 6 independent per group). (c) Experimental design of the study. SH-SY5Y or BV-2 cells were separately cultured in different plates. Vehicle or mdivi-1 (25 μM) was pre-treated in SH-SY5Y or BV-2 cells, then the media was changed to glucose-containing media (Ctrl, 25 mM) or glucose-deprived media (0 mM) and co-cultured. (d) Representative confocal images of SH-SY5Y cells with cleaved caspase-3 staining. Scale bars, 50 μm. (e) Quantification of cleaved caspase-3^+^ cells as a percentage of DAPI^+^ cells (*n* = 6 or 9 independent samples per group). (f) Experimental design of the study. SH-SY5Y cells were incubated in normal glucose-containing media (Ctrl, 25 mM) or glucose-deprived media (0 mM) with vehicle (GD) or minocycline (HM, 50 μM) for 24 hours. (g) Representative confocal images of SH-SY5Y cells with cleaved caspase-3 staining. Scale bars, 50 μm. (h) Quantification of cleaved caspase-3^+^ cells as a percentage of DAPI^+^ cells (*n* = 5 or 6 independent samples per group). (i) Representative confocal images of SH-SY5Y cells with Mitotracker staining. Scale bars, 10 μm. (j) Quantification of mitochondrial length (*n* = 5 or 6 independent samples per group). (k) Experimental design of the study. BV-2 cells were incubated in normal glucose-containing media (Ctrl, 25 mM) or glucose-deprived media (0 mM) with vehicle (GD), mdivi-1 (25 μM), or minocycline (50 μM) for 24 hours. (l) Representative confocal images of BV-2 cells with CD68 staining. Scale bars, 20 μm. (m) Quantification of CD68 intensity (*n* = 6 independent samples per group). Statistical analysis were performed using multiple t tests with Holm-Sidak method (a and b) and one-way ANOVA followed by Tukey’s multiple comparison test (e, h, j, and m). Data are presented as means ± SEM. **P < 0.01; ***P < 0.001.

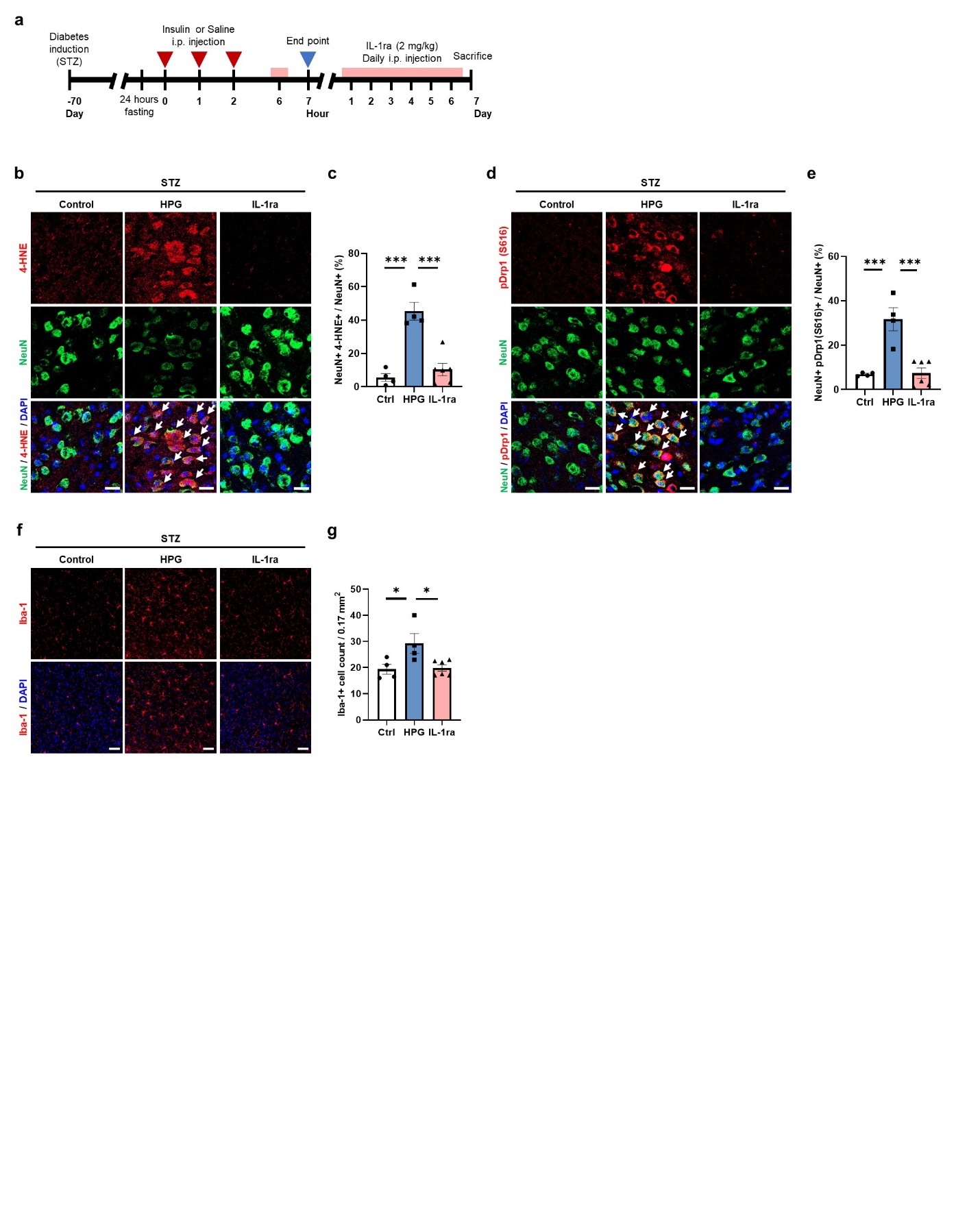

**Supplementary Fig. 5. IL-1 signaling blockade prevent hypoglycemia-induced neuronal damage in STZ-induced diabetic mice. Related to Figure 5.** (a) Experimental design of the study. All C57BL/6 mice were i.p. injected with STZ (55 mg/kg) for five consecutive days. After diabetes induction (blood glucose level exceeding 200 mg/dL), diabetes condition was maintained for 10 weeks prior to hypoglycemia experiment. The diabetic control group without insulin-induced hypoglycemia (Control; *n* = 4) received vehicle injection, whereas diabetic mice subjected to insulin-induced hypoglycemia were treated with either vehicle (HPG; *n* = 4) or IL-1ra (IL-1ra; *n* = 6). (b) Representative confocal images of the RSC tissues with NeuN and 4-HNE co-staining. Scale bars, 20 μm. White arrows indicate 4-HNE^+^ cells co-localized with NeuN^+^ cells. (c) Quantification of the percentage of 4-HNE^+^ cells co-localized with NeuN^+^ cells. (d) Representative confocal images of the RSC tissues with NeuN and pDrp1(S616) co-staining. Scale bars, 20 μm. White arrows indicate pDrp1^+^ cells co-localized with NeuN^+^ cells. (e) Quantification of the percentage of pDrp1^+^ cells co-localized with NeuN^+^ cells. (f) Representative confocal images of the RSC tissues with Iba-1 staining. Scale bars, 50 μm. (g) Quantification of Iba-1^+^ cells normalized by the area size of 0.17 mm^2^. Statistical analysis was performed using one-way ANOVA followed by Tukey’s multiple comparison test. Data are presented as means ± SEM. *P < 0.05; ***P < 0.001.

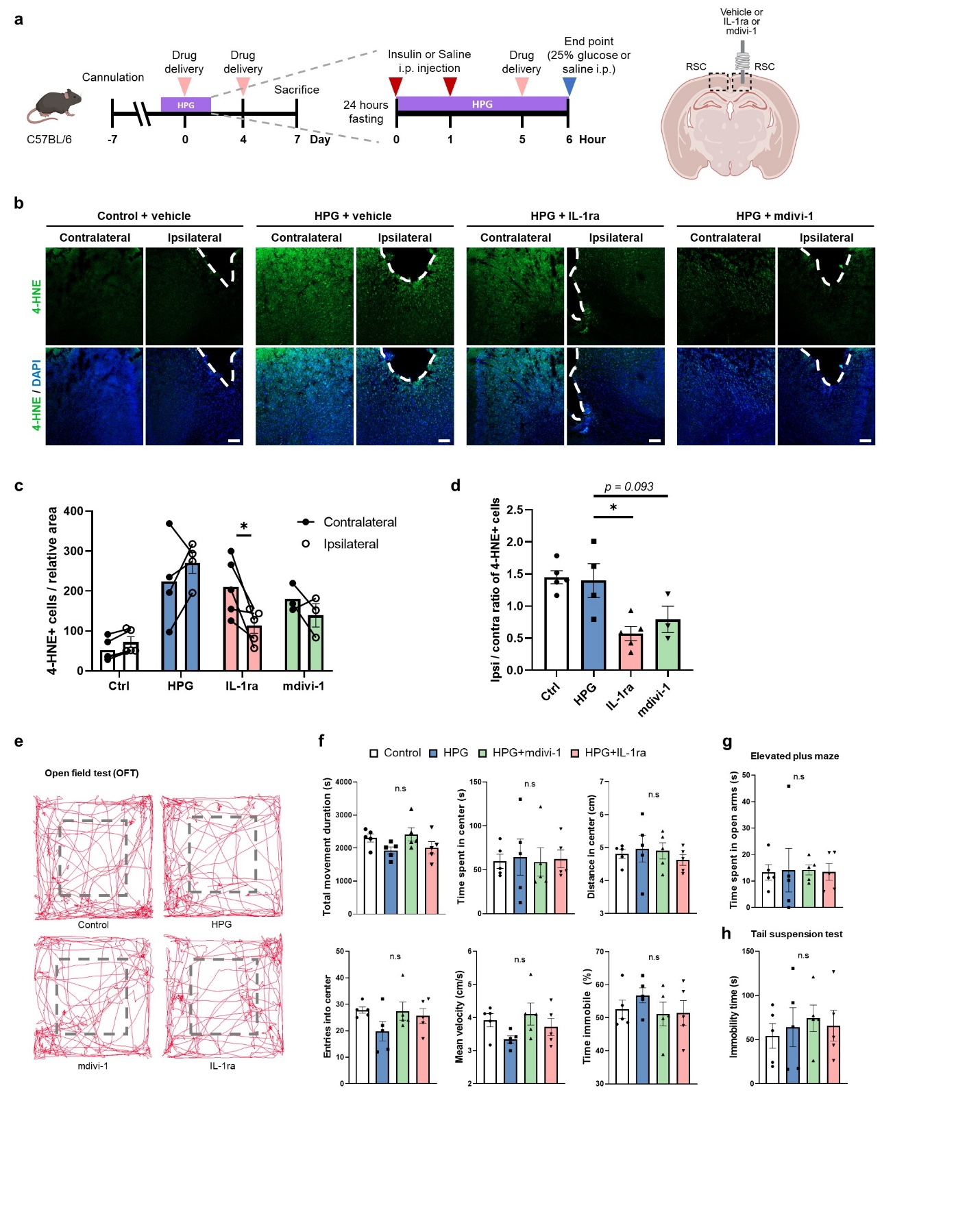

**Supplementary Fig. 6. RSC-specific inhibition of mitochondrial fission and IL-1 signaling, and additional behavioral characterization. Related to Figure 6 and 7.** (a) Experimental design of the study. (b) Representative confocal image of 4-HNE^+^ cells in the ipsilateral and contralateral sites of the RSC. Scale bars, 100 μm. (c) Quantification of 4-HNE^+^ cells in the relative area size of the contralateral and ipsilateral sites of the RSC tissue. (Connecting lines represent matched slices from the ipsilateral and contralateral sites of the RSC within the same mouse.) (d) The ratio of 4-HNE^+^ cells in the ipsilateral/contralateral sites in the RSC. (Ctrl, *n* = 5 mice; HPG, *n* = 4 mice; IL-1ra, *n* = 5 mice; mdivi-1, *n* = 3 mice). (e) Representative image of open field test. (f) Quantification of locomotor activity and anxiety-related behavior assessed through the open field test (*n* = 5 mice per group). (g) Quantification of time spent in the open arms during the elevated plus maze (*n* = 5 mice per group). (h) Quantification of immobility time during the tail suspension test (*n* = 5 mice per group). Statistical analysis were performed using multiple *t* tests with Holm-Sidak method (c) and one-way ANOVA followed by Dunnett’s multiple comparisons test (d, f, g and h). Data are presented as means ± SEM. *P < 0.05; ns, not significant.
